# The Genetic Landscape of Transcriptional Networks in a Combined Haploid/Diploid Plant System

**DOI:** 10.1101/007153

**Authors:** J.-P. Verta, C.R. Landry, J. MacKay

## Abstract

Heritable variation in gene expression is a source of evolutionary change and our understanding of the genetic basis of expression variation remains incomplete. Here, we dissected the genetic basis of transcriptional variation in a wild, outbreeding gymnosperm (*Picea glauca*) according to linked and unlinked genetic variants, their allele-specific (*cis*) and allele non-specific (*trans*) effects, and their phenotypic additivity. We used a novel plant system that is based on the analysis of segregating alleles of a single self-fertilized plant in haploid and diploid seed tissues. We measured transcript abundance and identified transcribed SNPs in 66 seeds with RNA-seq. Linked and unlinked genetic effects that influenced expression levels were abundant in the haploid megagametophyte tissue, influencing 48% and 38% of analyzed genes, respectively. Analysis of these effects in diploid embryos revealed that while distant effects were acting in *trans* consistent with their hypothesized diffusible nature, local effects were associated with a complex mix of *cis*, *trans* and compensatory effects. Most *cis* effects were additive irrespective of their effect sizes, consistent with a hypothesis that they represent rate-limiting factors in transcript accumulation. We show that *trans* effects fulfilled a key prediction of Wright s physiological theory, in which variants with small effects tend to be additive and those with large effects tend to be dominant/recessive. Our haploid/diploid approach allows a comprehensive genetic dissection of expression variation and can be applied to a large number of wild plant species.

## Introduction

Evolutionary changes are fueled by genetic variation that translates to molecular, cellular and physiological levels, upon which natural selection acts. Gene transcription plays a pivotal role in the transfer of genetic variation to phenotypes as it is the first functional step from information encoded in the genome (Fay and Wittkopp, 2008, Emerson and Li, 2010, Romero *et al.*, 2012, Wittkopp and Kalay, 2012). Variation in transcript abundance correlates with variation in protein translation (Muzzey *et al.*, 2014) and abundance (Albert *et al.*, 2014), and translates into cellular phenotypes such as human cancer (Gregory and Cheung, 2014), variation in physiological traits such as chemical composition of the cell wall in trees (MacKay *et al.*, 1997) and in morphological phenotypes such as defensive pelvic apparatus in fish (Chan *et al.*, 2010), for example. Given the critical role that gene transcription plays in determining phenotypes, investigating the genetic bases of expression variation is central to our understanding of evolution.

Dissection of the genetic architecture of expression variation will help us to uncover the driving forces behind its emergence, persistence and divergence. As for any quantitative trait, determining the genetic architecture of expression variation includes identifying the number of loci involved and their genomic positions, the molecular mechanisms by which such loci influence expression and their homozygous and heterozygous effects on the expression phenotype (Mackay, 2001). The first of these goals can be achieved by mapping the positions of genetic effects influencing transcription (expression). For any given gene (the focal gene), genetic effects typically map to multiple linked and unlinked loci (Brem et al., 2002), and are here referred to as local and distant effects, respectively. The molecular mechanisms by which local and distant effects influence expression can be inferred from the allele-specificity of these effects. Genetic effects representing diffusible molecules are expected to influence the expression of both alleles of a diploid cell in a similar fashion, while non-diffusible effects are likely to be allele-specific (Wittkopp *et al.*, 2004). Allele-specific and non-specific effects are here referred to as *cis* and *trans*, respectively (Landry *et al.* 2005, Rockman and Kruglyak, 2006, Ronald *et al.*, 2005, Skelly *et al.*, 2009, Emerson and Li, 2010). In general, unlinked effects are assumed to influence focal gene expression levels in *trans* through diffusible molecular signals, such as protein or RNA (Wittkopp *et al.*, 2004, Tirosh *et al.*, 2009, McManus *et al.*, 2010, Emerson *et al.*, 2010, Emerson and Li, 2010, Gruber *et al.*, 2012, Goncalves *et al.*, 2012, Meiklejohn *et al.*, 2013, Coolon *et al.*, 2014). Genetic effects that are linked with the focal gene are often interpreted as local sequence variants that alter the affinity of transcriptional regulators, mRNA stability or chromatin states in *cis* (Goncalves *et al.*, 2012, Wittkopp and Kalay, 2012, Chang *et al.*, 2013, Kilpinen *et al.*, 2013, McVicker *et al.*, 2013). Considerable deviation is however expected from these predictions, as local variants can act in *trans* through diffusible molecules (e.g. Jacob and Monod, 1961, Ronald *et al.*, 2005) and distant effects in *cis* through direct interactions between chromosomes (e.g. Spilianakis *et al.*, 2005, Kilpinen *et al.*, 2013, Jin *et al.*, 2013, Battle *et al.*, 2014).

Persistence of expression variation within and expression divergence among populations depends on the efficiency of natural selection to purge detrimental and to select for beneficial variation. Genetic variation with beneficial additive or dominant phenotypes are thought to be a likely source of evolutionary change because they are exposed to natural selection as soon as they emerge in a population and can thus quickly reach fixation (Hartl *et al.*, 1997). Time for new recessive beneficial mutations to reach fixation is significantly longer, and such alleles are expected to contribute relatively little to adaptive differences (i.e. Haldane s sieve, Haldane, 1927). In addition, even a low level of recessivity allows harmful variation to persist in populations at low frequencies under mutation selection balance (Hartl *et al.*, 1997). Distinguishing between additive and dominant/recessive modes of inheritance therefore becomes a key in understanding the evolutionary dynamics of expression variation in cases such as outbreeding species where heterozygosity is common (McManus *et al.*, 2010, Lemos *et al.*, 2008).

The relationships between the location, allele specificity and phenotypic additivity of genetic effects on expression are a key in determining the genetic bases of expression variation (Rockman and Kruglyak, 2006). Haploid/diploid systems are a powerful tool for this task (Ronald *et al.*, 2005). The haploid phase of these systems facilitate discovery of genetic effects because dominance is absent and effects should segregate 1:1. The allele-specificity and dominance relationships of the effects can be thereafter tested in diploids. Studies leveraging on a haploid/diploid approach have been relatively rare, and studies of the genetic architecture of expression variation have mostly investigated interspecific crosses (e.g. McManus *et al.*, 2010), or predominantly inbreeding species (e.g. Zhang *et al.*, 2011) where heterozygosity is limited. Other studies have considered new mutations instead of standing variation (Gruber *et al.*, 2012), or excluded the possibility of self-regulating (local *trans*) effects (Meiklejohn *et al.*, 2013). Overall, very few documented systems exist that permit the comparison of effect linkage, allele-specificity and inheritance, and even fewer are the systems where this can be applied directly to individuals from outbreeding natural populations.

We recently explored an outbreeding system in a multicellular eukaryote where genetic effects on gene expression can be identified in a haploid state (Verta *et al.*, 2013). This system is based on measuring gene expression in the megagametophyte tissue of the seed of gymnosperm plants. Megagametophytes are haploid products of meiosis and their genomes are maternally inherited (Fig. 1A, Williams, 2009). Here, we expand this biological system to include a diploid segregating tissue by self-fertilizing a white spruce (*Picea glauca* [Moench. Voss]) individual. In a self-cross, alleles of loci that are heterozygous in the mother tree segregate 1:1 in the megagametophytes and in 1:2:1 genotype combinations in the embryos (Fig. 1B). This enables measurement of the impacts of the same genetic variants in haploid and diploid tissues. Another advantage of the system is that both the megagametophyte and embryo inherit identical maternal genomes (Fig. 1B), facilitating genetic tracking of the progeny. We use this haploid/diploid system in conjunction with RNA-seq (Fig. 1C) to evaluate the intersect between the genomic location and allele specificity of genetic effects on gene expression as well as their additive versus dominant/recessive mode of inheritance (Fig. 1D).

**Figure 1.**
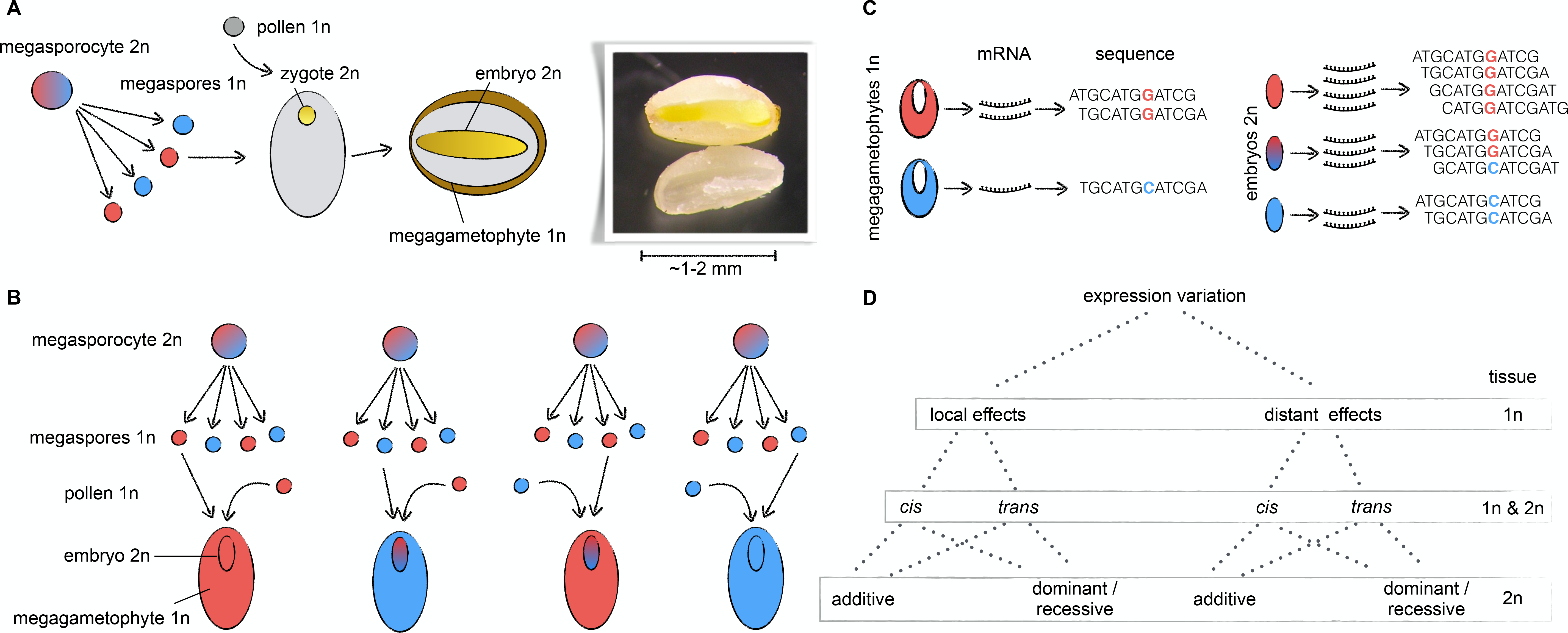
Spruce haploid/diploid system. **A)** Seed development. Red and blue colors represent the two alleles at a given locus. **B)** Segregation of alleles in megagametophytes and embryos in a self-cross where the pollen is also the meiotic product from the mother tree. There are two possible haploid genotypes (megagametophytes) and three possible diploid genotypes (embryos) for each heterozygous locus in the mother’s genome. **C)** RNA-seq in haploids and diploids facilitates the measurement of allele expression levels (number of reads is proportional to expression level of the allele) and transcribed SNPs. **D)** Genetic dissection of expression variation indicating tissue (1n or 2n) used for each analysis step in the study.

## Results

### Haploid comparison

We performed transcriptome sequencing (RNA-seq) of 66 megagametophytes originating from a single wild mother tree. For simplicity, we will refer to megagametophytes as haploids in this report. Reads were aligned to white spruce transcribed gene models (representing unique genes, Rigault *et al.*, 2011, Table S1). The average portion of uniquely aligned reads and the overall alignment rate were respectively ∼50% and ∼70%, on average. Comparison of read counts in samples that were replicated within and between sequencing lanes indicated that reproducibility was extremely high (Pearson’s correlation 0.999 and 0.998, respectively), and that variation in read counts due to biological variation among samples was much larger than variation assigned to technical sources (Fig. S1). We detected expression (defined as at least 90% of the samples having at least one read for a given gene) of 15,736 of the 27,720 genes represented in the white spruce gene catalogue (Rigault *et al.*, 2011). The number of expressed genes was in line with earlier studies based on microarrays that identified the megagametophyte as one of the spruce tissues with the largest number of expressed genes (Raherison *et al.*, 2012).

We first tested whether genetic sources of expression variation were in close genetic linkage with the expressed genes. We examined this by testing for expression differences between allele classes for each gene, identified based on SNPs in the RNA-seq reads. Sequence variation was observed in 6257 genes, each with one (28%) or multiple (72%) SNPs that segregated following Mendelian 1:1 expectations in the haploids (Fig. S2). Base calls on SNP positions were 96% identical across all samples. We assigned each haploid sample to one of the two alleles of each gene and compared the numbers of RNA-seq reads assigned to allele classes with a likelihood ratio test. Haploid samples were considered as biological replicates for each allelic class in this approach. Abundant local variation in gene expression was observed, influencing roughly half (48%) of the genes with SNPs (*q*<0.01, Table S2). These genes were assigned as being under local effects. This abundant local genetic variation indicates that genetic variants in the genomic proximity of focal genes have a large-scale influence on expression levels.

Next, we examined cases where the genetic bases of expression variation were unlinked to the focal gene. We tested for association between the expression levels of all genes and genotypes of unlinked loci inferred from RNA-seq data (94,245,480 tests). Genes were grouped based on linkage between genotypes into 12 linkage blocks of nearly equal size, consistent with the haploid chromosome number of spruce, and a few smaller blocks (Fig. S3). Genes were left unordered within these blocks. Focal genes not containing SNPs could not be mapped, and any association with markers in other loci could not be clearly determined as local or distant. We therefore proceeded with independent analyses of the distant associations for these genes.

We observed that 38% of tested focal genes were influenced by distant effects (*q*<0.01, Table S2). Most of the distant effects were observed in a number of loci assigned to the same linkage block, which were counted as a single effect (seen as horizontal lines within linkage blocks in Fig. 2). Overall, distant effects were distributed randomly over linkage blocks (Supplemental text). Their relatively high abundance compared to local effects, which are more likely to be discovered due to a higher statistical power, points towards a complex and highly connected control of gene expression levels across linkage blocks.

**Figure 2.**
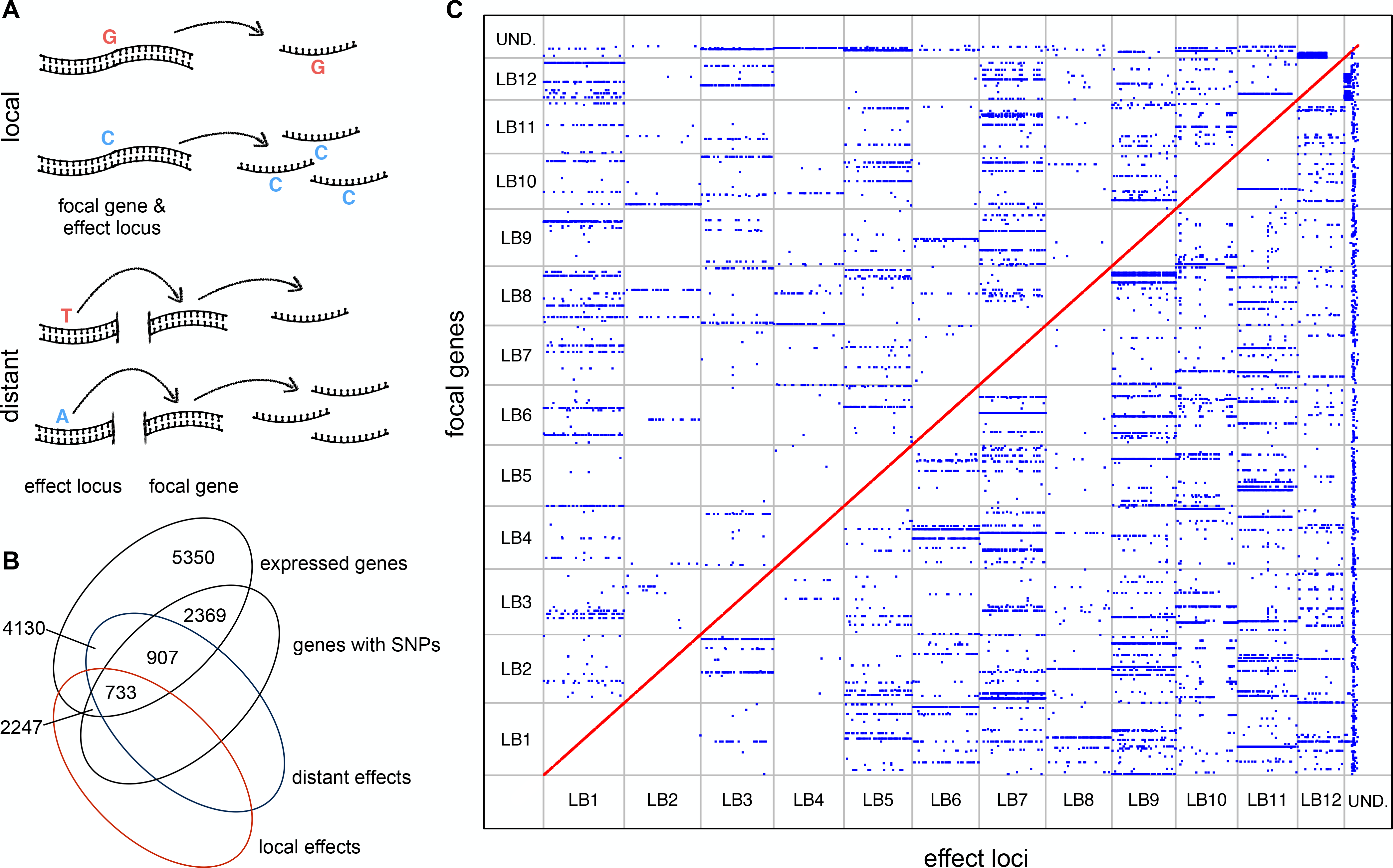
Mapping local and distant effects. **A)** Local effects were defined as genetically linked to the expressed gene, while distant effects were unlinked. **B)** Distribution of focal genes, according to detected SNPs and genetic effects. **C)** Position of local and distant effect (red and blue points, respectively) loci versus positions of the focal gene. For distant effects, only associations that involve mapped focal genes are shown. Twelve largest linkage blocks (LB1-LB12) are delimited with gray lines. Data points outside these represent smaller linkage blocks and focal genes not assigned to linkage blocks (UND).

We compared the local and distant genetic effects to the cases where our previous investigation of the same individual tree indicated that expression variation in the focal gene was associated with a single major effects locus (Verta *et al.*, 2013). Here we analyzed 397 of the genes under major genetic effects, of which one hundred exhibited local effects and 262 exhibited distant effects. Taking both local and distant effects into account, 91% of the major genetic effects observed in our previous study could be identified here, i.e. over sixty times more than expected by chance alone. The majority of local and distant effects identified in this report explained expression variation in the progeny only partially and likely represented parts of a polygenic basis of expression variation (Supplemental text and Fig. S4).

### Haploid/diploid comparison

We next examined whether the genetic effects identified in the haploids translated into genetic effects in the diploids (embryos). RNA-seq for 66 self-fertilized embryos were obtained from the same seeds used for haploid analysis (Fig. 1) and aligned to white spruce gene models (Rigault *et al.*, 2011, Table S1), producing similarly high alignment rates. The expression of 15,859 genes was detected in the diploids, indicating that the embryos were at least as transcriptionally active as the megagametophytes.

Genetic effects on gene expression can be specific to diploid tissues (e.g. Drost *et al.*, 2010). We therefore separated genes into preferentially expressed in the haploids, diploids, or non-preferentially expressed. Most expression differences between tissues were of small magnitude; average expression levels in haploids were highly correlated with expression in diploids (Figs. 3A and S1L). A total of 2853 genes that varied by log_2_ fold change of two or higher were deemed as tissue preferential. The majority of genes were assigned as non-preferentially expressed, indicating that expression divergence was relatively minor and allow genetic effects to be compared across haploids and diploids (Figs. 3A and S1L).

**Figure 3.**
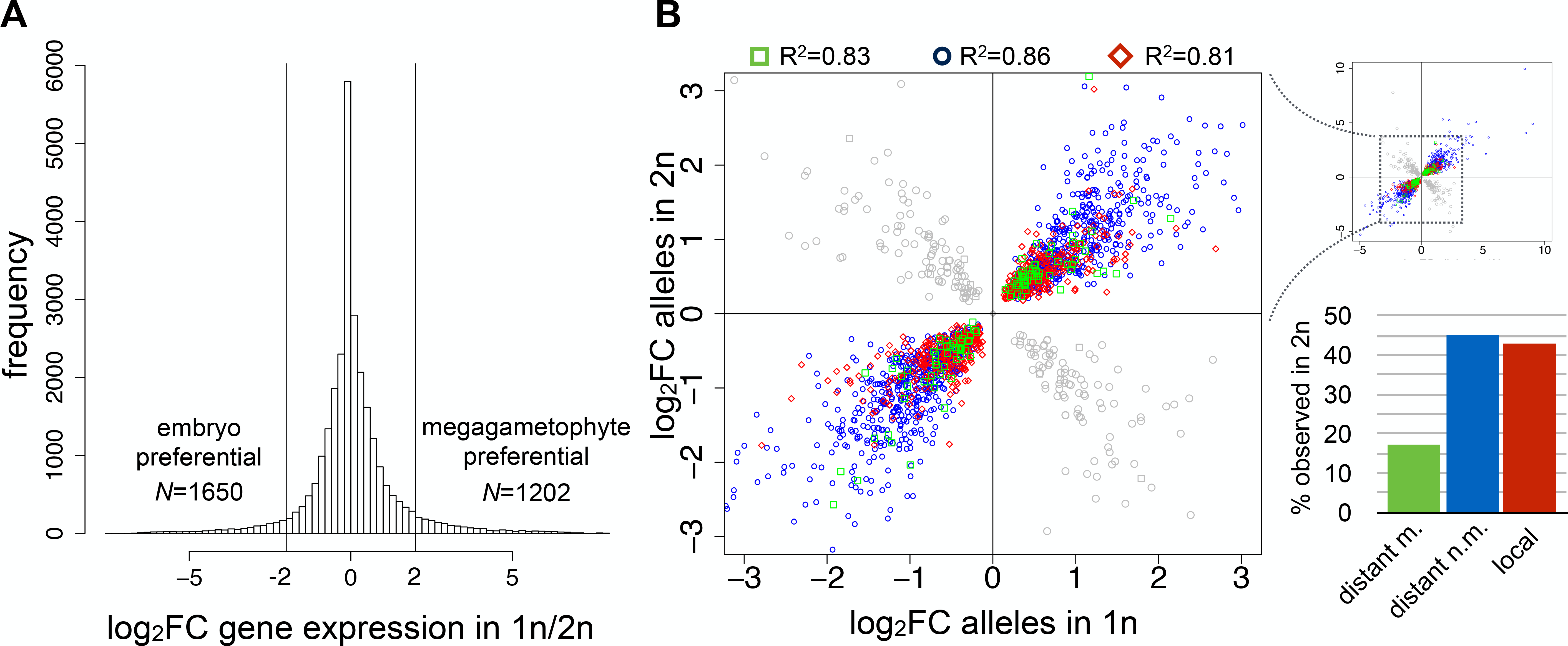
Genetic effects on gene expression in haploids and diploids. **A)** Tissue-preferential expression between haploid and diploid tissues. Log_2_FC of two or more was used to designate tissue-preferential expression (19.5% of genes). **B)** Effects that were observed between homozygous diploids. Log fold change of expression levels between allele classes are plotted in haploids (1n x-axis) against that in diploids that were homozygous for the given effect (2n y-axis). Bars (right) represent the proportion of tested local and distant effects (discovered in haploids) that were observed between diploids homozygous for the effects (see Tables S3 and S4). Coefficient of determination (R^2^) of linear models (expression in diploids as dependent variable and expression in haploids as linear predictor) for each effects category are given above the scatterplot. Distant m. (green squares); distant effects influencing mapped focal genes, distant n.m. (blue circles); distant effects influencing non-mapped focal genes, local (red angled squares), grey symbols; effects that change signs between haploids and diploids and that were excluded from downstream analysis.

As a first step to compare genetic effects in haploids and diploids, we investigated whether local and distant effects could be observed in diploid homozygotes. We tested for differential expression between diploid genotype classes that were homozygous for the local and distant effects. Proportions of effects that were recapitulated between homozygous diploids ranged from 17% to 45% (*q*<0.01, Fig. 3B and Tables S3 and S4). Effects that explained a larger part of expression variation were more likely to be recapitulated in homozygotes (t-test *P*<2.2e-16). The identity of high and low expression alleles remained largely unchanged (82% for local effects and 86% for distant effects, Fig. 3B). In a small proportion of the cases (∼15 %) the relative expression levels of the alleles were reversed in the diploids, a phenomenon that has been observed in haploid/diploid yeast (Ronald *et al.*, 2005, Gruber *et al.*, 2012). Inclusion of these effects in downstream analyses did not change our conclusions, but we nonetheless excluded these cases of expression reversals because they may result from complex genetic-ploidy interactions. The set of effects that were observed in homozygotes forms an important reference point in our further analyses where effects are compared across diploid genotypes.

### Allele specific expression

Having established a set of genetic effects observed across ploidies, we focused on heterozygous diploids to examine the allele-specificity (*cis* versus *trans*) of local and distant effects. The distinction between *cis* and *trans* can be established by testing whether a genetic variant influences the expression of one, or two alleles of the focal gene in heterozygous individuals (Wittkopp *et al.*, 2004). We first tested for *cis* effects between the two alleles in heterozygous diploids in order to establish a set of genes under *cis* effects to which haploid expression would be compared. A third (31%) of tested genes showed expression differences between the alleles (*q*<0.01, Table S3), indicating that *cis* effects were common in diploids. Next, we investigated the relationship between locally linked variants and allele-specific (*cis*) effects. Technical differences in determining effects in haploids and diploids were taken into account by defining a set of local effects that were observable based on SNP counts (Methods and Supplemental text). We observed that 55% of the genes under local effects were also associated with *cis* effects (over four and half times the frequency expected by chance), while 75% of the genes that showed *cis* effects were assigned as under local effects (over six times the frequency expected by chance, Fig. 4A, 4B and 4C, Table S3). Relative allele expression levels remained unchanged in the large majority of the cases (96%, Fig. S6). These results suggested that allele-specific effects were more often due to local genetic variants than the contrary, where local genetic effects were acting in an allele-specific manner.

**Figure 4.**
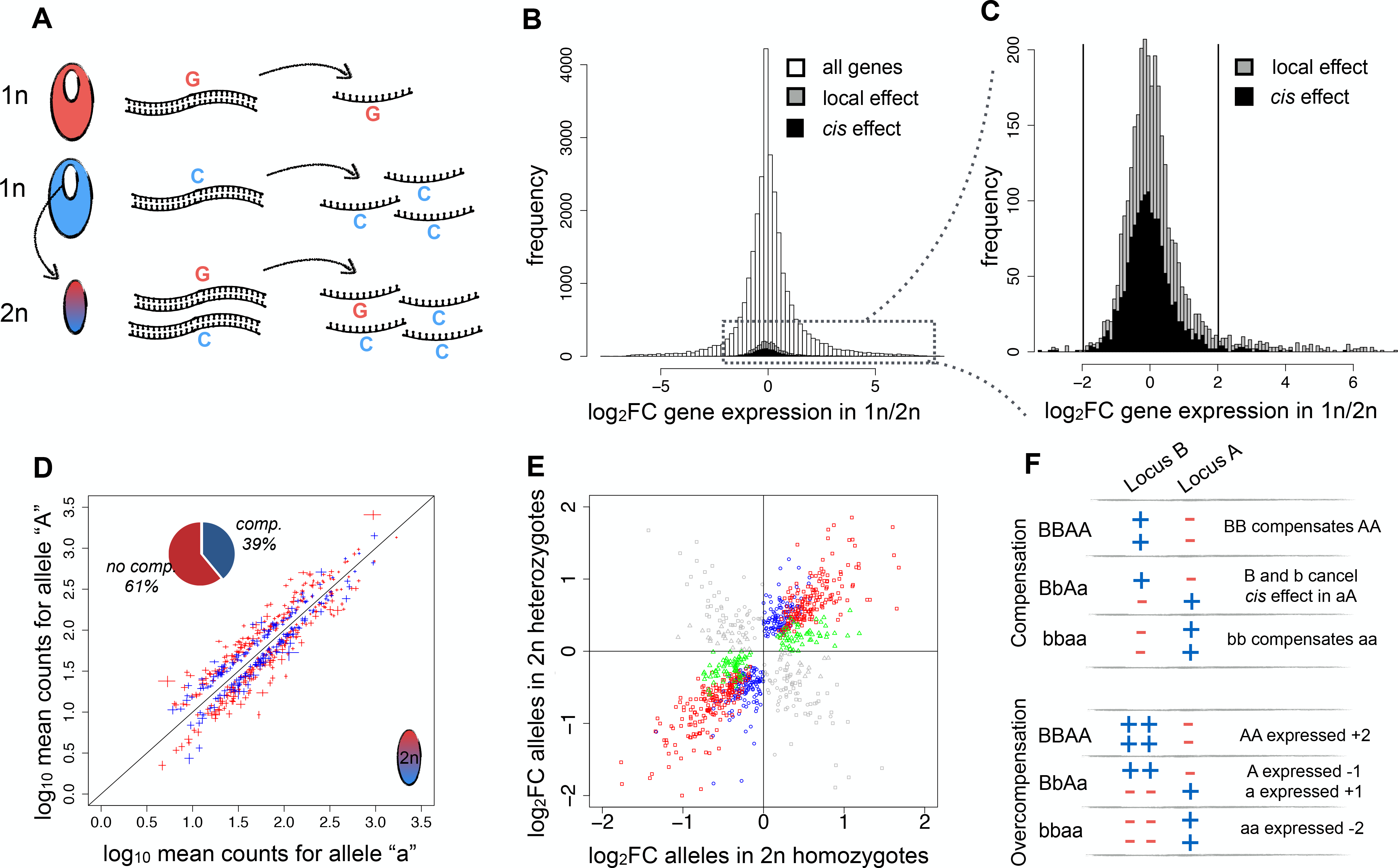
Local effects in *cis* and *trans*. **A)** Allelic expression differences due to local effects observed in haploids were compared to allele expression differences in heterozygous diploids. **B)** Tissue preferential expression levels in all genes (white), genes under local effects in haploids (grey) and genes under *cis* effects in diploids (black). **C)** Close-up of tissue-preferential expression in genes under local (grey) and *cis* effects (black). Threshold of tissue-preferential expression correspond to log_2_FC values −2 and 2. **D)** Comparison of allele counts within heterozygous diploids. Each data point represents a gene under local *cis* effect. Horizontal and vertical lines intersect at the mean value of allele expression levels and show the standard error of the mean counts of each allele. Blue: genes under compensatory effects, red: genes where no compensation was observed. Notice there is no difference in the distribution of blue and red data points and the standard errors never cross the diagonal line, which marks equal expression of the two alleles. Pie-chart, proportions of local *cis* effects with expression differences between homozygotes (no compensation, no comp.) and those where no homozygote difference was observed (compensation, comp.). **E)** Allele expression levels (log_2_FC) between homozygous diploids (x-axis) and within heterozygous diploids (y-axis). Genes under local *cis* effects with and without compensation are represented as blue circles and red squares, respectively. Local *trans* effects, green triangles. Grey symbols correspond to effects that change signs between homozygotes and heterozygotes (overcompensation) and that were excluded from downstream analyses. Including these cases in downstream analyses did not change the conclusions. **F)** Schematic of compensation and overcompensation. Locus A represent the focal genes the expression of which is being measured and locus B the compensatory effect in linkage. Plus and minus signs represent signs of effects on focal gene expression.

Comparisons across diploid genotypes were used to identify local genetic variation that acted in *trans*. The expectation for *trans* effects was that homozygous diploids would have different expression levels, but no differential expression between alleles would be detected in heterozygotes (Wittkopp *et al.*, 2004, Fig. 4E). Following this reasoning, any homozygote difference in genes under local effects that were not associated with *cis* effects were designated as local effects acting in *trans*. These were observed in 12% of the tested genes (Table S3). Having equal effects on both of the alleles in heterozygotes, these effects were candidate diffusible signals that were due to self-regulation, or independent genes situated in close linkage to the expressed gene.

Comparison across homo-and heterozygote diploids facilitated further dissection of local effects. We first investigated whether local *cis* effects could be observed in homozygous diploids in addition to the heterozygotes. The majority (438 focal genes, 61% of tested) of local *cis* effects were also detected in homozygotes (Fig. 4D and 4E), while in 284 focal genes (39% of tested) *cis* differences were observed in heterozygotes only (Fig. 4E, 4F and S7). We verified that allelic expression differences were consistent between individual heterozygous diploids, i.e. the same allele was more strongly expressed in all heterozygotes (Fig. 4D). Previous studies in interspecific hybrids have reported *cis* effects specific to heterozygous genotypes, and interpreted such patterns as compensation of the *cis* effect by different *trans* backgrounds in the homozygotes (e.g. Wittkopp *et al.*, 2004, Landry *et al.*, 2005, McManus *et al.*, 2010). The observed patterns of heterozygote-specific *cis* effects were consistent across the progeny, i.e. largely independent of the genetic background that segregates separately from the focal gene. This suggests that either the compensatory factors were in very close linkage to the focal gene, or that these effects were caused by different mechanisms, such as interaction between the *cis* effect alleles, which we find unlikely. Finally, 165 focal genes under either of local *cis* or local *trans* effects changed signs between homozygotes and heterozygotes (18% of tested, Fig. 4E). This pattern is also compatible with either compensation by a closely linked factor the effect of which is larger than that of the focal effects (termed here as “overcompensation”, Fig. 4F, Goncalves *et al.*, 2012), or by an interaction between the *cis* effect alleles. Including these effects in downstream analyses did not change the conclusions. We however excluded the cases where local effects changed signs in order for our analysis to be linear across genotypes.

We determined whether distant effects identified in the haploids were associated with *cis* or *trans* effects in the diploids. Distant effects on homozygous genes were assumed to act in *trans* because genes under *cis* effects are expected to be polymorphic (Zhang and Borevitz, 2009, McManus *et al.*, 2010, Zhang *et al.*, 2011, Goncalves *et al.*, 2012), yet no *cis*-divergence in the focal gene RNA-seq reads could be observed. The hypothesis that distant variants act in *cis* was testable in embryos that were heterozygous for both the distant and the focal genes. Patterns consistent with distant *cis* effects were discovered in a frequency corresponding to our false discovery rate (45 of 3665 tested associations) and the segregating frequencies of these indicated that the association pairs were in fact linked in most cases. The overwhelming majority of distant effects therefore acted in *trans*, hence likely representing diffusible signals.

### Additive versus dominant/recessive mode of inheritance of expression variation

Our third objective was to investigate the additive versus dominant/recessive mode of inheritance of genetic effects. We compared focal gene expression levels between diploids that were homo- and heterozygous for the genotypes associated with the genetic effects. For additive effects, statistically significant expression differences were expected between heterozygotes and both of the homozygotes (Fig. 5). In contrast, dominance was expected to produce a statistical difference between the heterozygous genotype and only one of the homozygotes (the recessive genotype).

**Figure 5.**
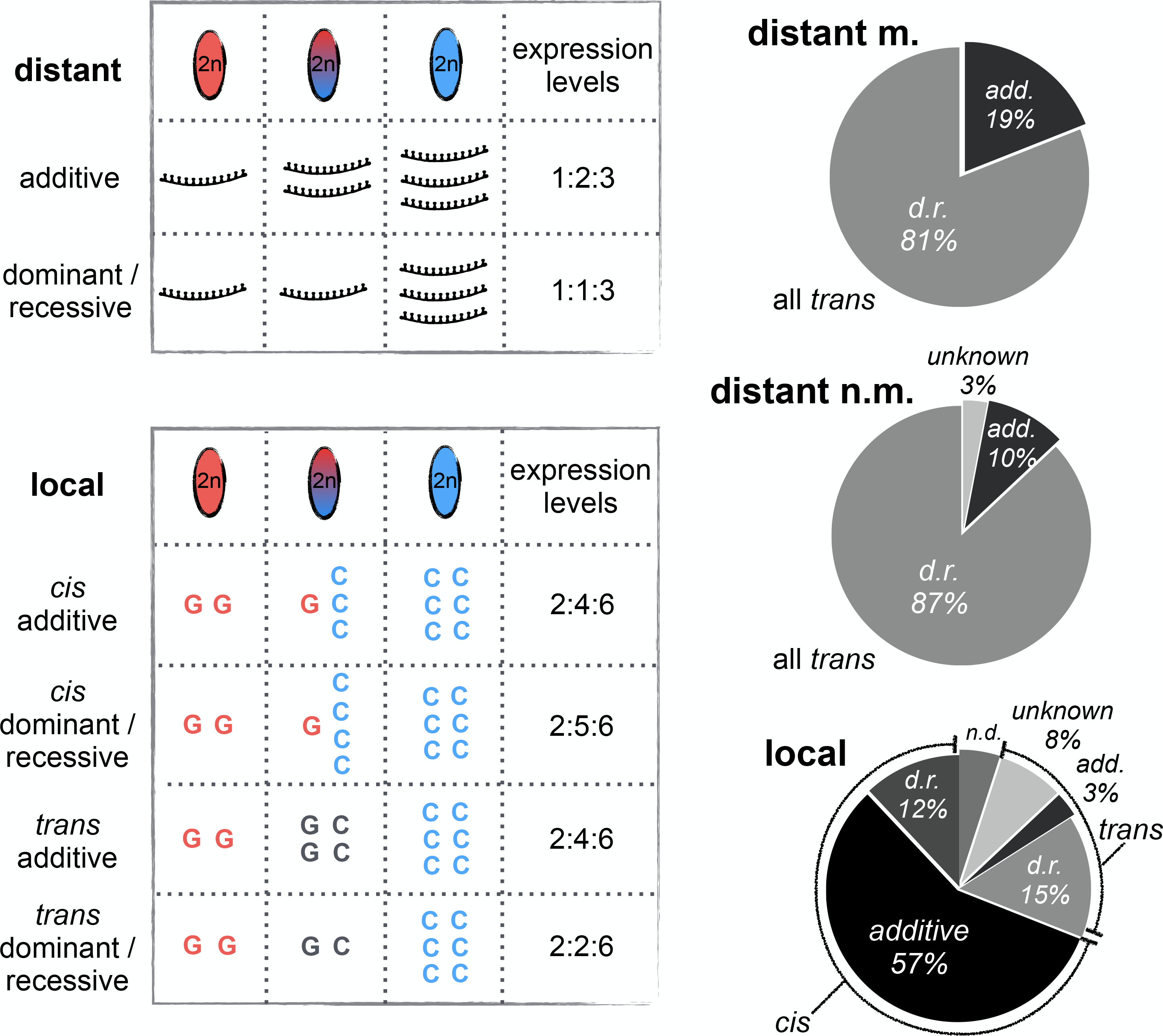
Additive versus dominant/recessive modes of inheritance of local and distant effects. Cartoon tables (left) of different possible categories of allele specificity and inheritance. Only cases of complete dominance (gene expression in heterozygote is identical to homozygous dominant) and additivity (gene expression in heterozygote is exactly at midpoint between homozygotes) are shown for clarity. mRNA illustrations (distant effects) and letters corresponding to nucleotides (local effects) depict allelic expression levels in different embryo genotypes. Pie-charts (right) show the proportions of local and distant effects observed in haploid and all diploid genotypes under each category. Proportions of each effect category are calculated from tables S3 and S4. For local effects, the relative proportions of *cis* and *trans* observable between all three genotype classes of diploids are calculated as ratio of column 4 on column 5 of table S3. Proportions of different modes of inheritance within *cis* and *trans* effects are calculated from percentages in columns 6-9 of table S3. Relative frequencies of local *cis* and *trans* effects are different from columns 3 and 4 of table S3 because some local *cis* effects were observed only in heterozygotes and did not correspond to differences between homozygous diploid genotypes. “Distant m.” and “distant n.m.” effects represent non-redundant effects that influence mapped and non-mapped focal genes, respectively. d.r.; dominant/recessive, add.; additive, n.d.; non-determined.

Different patterns of inheritance were observed for effects acting in *cis* and *trans*. Expression phenotypes were mostly additive in genes under local *cis* effects (83% additive vs. 17% dominant/recessive, *q*<0.01), whereas those under local *trans* effects were more dominant/recessive (59% dominant/recessive vs. 10% additive, 31% unknown, *q*<0.01, Table S3 and Fig. 5). Expression variation followed a dominant/recessive inheritance in the large majority of distant effects as well, all of which acted in *trans* (*q*<0.01, Table S4 and Fig. 5). The observed clear distinction in the inheritance of local *cis* and *trans* effects suggested that the allele-specificity of the effects was important in determining their level of additivity.

We next investigated the level of phenotypic masking by dominance relationships by calculating a ratio between the midpoint expression level separating homozygotes and the difference of the heterozygote expression level from this midpoint (D/A ratio, Gibson *et al.*, 2004, Fig. 6A). As expected, D/A ratios largely recapitulated the identity of additive and dominant/recessive effects in each category (Fig. 6B). In dominant/recessive effects, a relatively low difference between heterozygotes and the homozygote midpoint (absolute D/A values in majority between zero and one, Fig. 6B) suggested that phenotypic masking by dominance relationships was not complete. Clearest distinction between dominance and additivity in expression was observed between distant *trans* effects influencing non-mapped focal genes and local *cis* effects, likely because these represented the classes with most observations (Fig. 6B).

**Figure 6.**
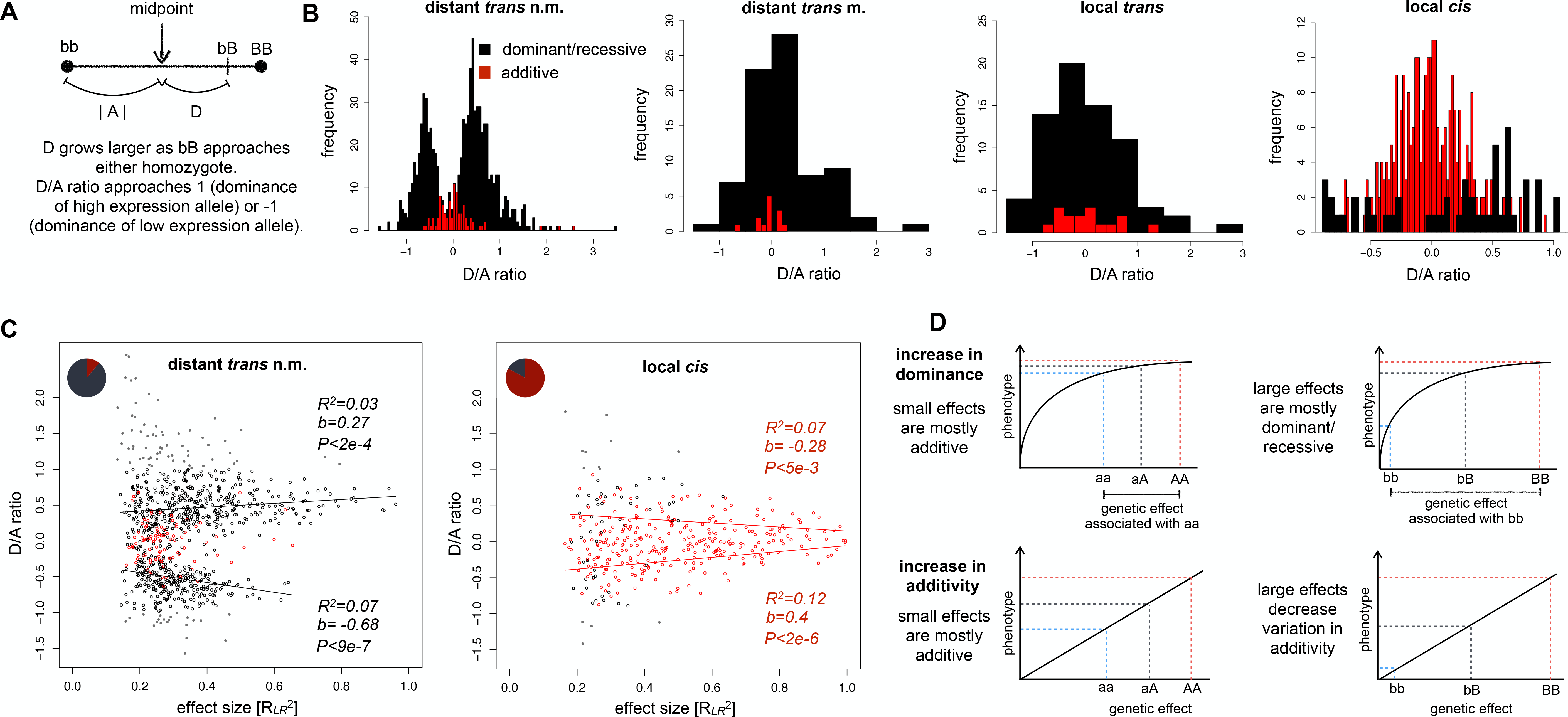
Phenotypic masking of heterozygote expression by dominance relationships and their correlation with effect size. **A)** D/A ratio. Distance between genotypes (bb, bB and BB) represents expression difference. D/A ratios close to zero indicate that the mean of heterozygote expression is halfway between mean homozygote expression levels (complete additivity) while ratios close to 1 or −1 indicate complete phenotypic dominance by one allele. Ratios beyond 1 and −1 indicate over- and under dominance, respectively. **B)** Distribution of D/A ratio in different effects. Dominant/recessive and additive associations are represented as black and red bars, respectively. **C)** D/A ratio of dominant/recessive (black open circles) and additive (red open circles) expression phenotypes (y-axis) is correlated with the strength of association (R*_LR_*^2^), which we use here as a relative measure of the proportion of explained variation by different genetic effects (x-axis, see Supplemental text and Methods, Rockman and Kruglyak, 2006). Grey points represent over- and underdominant effects which are not included in the theoretical predictions and therefore excluded from regression analysis. Pie-charts represent the proportions of dominant/recessive and additive effects in each category. Lines represent linear model fit between D/A ratio and strength of association for each category. “*b*” represents the slope of the linear regression model and describes the tendency of D/A ratio to increase with increasingly large effects in distant *trans* effects and to decrease in local *cis* effects. **D)** The physiological theory of dominance posits that recessivity of phenotypes is a consequence of a diminishing returns relationship between genotype and phenotype (Wright, 1934). According to Wright (1934), the hyperbolic response curve between genotype and phenotype is due to the fact that the individual gene product is not the rate-limiting factor in a given reaction. Kacser and Burns (1981) later showed that enzymatic reactions that involve multiple steps are robust to changes in the activity of the constituent factors, which is at the origin of the hyperbolic response curve. Mutations in factors that have larger homozygote differences correspond to more recessive phenotypes as the response curve between genotype and phenotype falls off the near linear portion, as was observed for *trans* effects. *Cis* effects were relatively robust to changes in effect size, which was consistent with a linear response curve with one rate-limiting factor (Kacser and Burns, 1981). Cartoon adapted from Wright (1934) and Phadnis and Fry (2005).

Next, we examined the relationship between homozygous genetic effect size (see Methods and Supplementary text) and the level of phenotypic masking by dominance. Wright’s physiological theory of dominance predicts that if the phenotype is the result of many interdependent steps, phenotypic masking by dominance should become stronger in variants associated with larger effects (Wright, 1934, Kacser and Burns, 1981, Phadnis and Fry, 2005). Overall, expression variation seems to follow predictions of the physiological theory of dominance: Distant *trans* effects that explained a larger proportion of expression variation were associated with D/A ratios closer to 1 or −1 (*P*<2e-4, Fig. 6C), indicating that phenotypic masking by dominance grew stronger with larger effects. In addition, distant *trans* effects classified as additive typically explained a small portion of expression variation, while effects classified as dominant/recessive typically had larger effects (seen as a gradient of red and black data points in Fig. 6C). Some effects of small size were classified as dominant/recessive, suggesting that a minority of effects may diverge from the predictions of Wright’s physiological theory (see: Kondrashov and Koonin, 2004, Agrawal and Whitlock, 2011). Interestingly, correlation between D/A ratio and effect size was nearly symmetric for D/A ratio values below and above zero. This indicates that phenotypic response to genetic variation is similar irrespective of whether heterozygote expression was closer to high or low expression homozygote (the phenotype response curve in Fig. 6D would be inverted for negative D/A ratios).

Wright’s theory also predicts that if the phenotype depends on a single rate-limiting factor, the level of dominance should be independent on the effect size or negatively correlated due to increased statistical power in larger effects (Fig. 6C). Such result was observed for *cis* effects; increasing effect size corresponded to stronger additivity of phenotypes (*P*<3e-3, Fig. 6C). Also local *trans* effects showed a significant correlation of stronger additivity with larger effects (*P*=3e-3, Fig. S8). These results are most consistent with a linear response curve between genotype and phenotype with one rate-limiting factor determining the phenotype (Wright, 1934, Kacser and Burns, 1981, Fig. 6C).

## Discussion

We dissected the genetic bases of expression variation in segregating haploid and diploid progeny of an outbreeding, wild gymnosperm plant. We analyzed the intersect between linked/unlinked genetic effects on gene expression and their allele-specific (*cis*) versus allele non-specific effects (*trans*), and additive versus dominant/recessive mode of inheritance in heterozygotes. We discovered abundant genetic variation in gene expression associated with local (linked) genetic variants (48% of genes) as well as distant (unlinked) variants (38%). The sets of genes under local and distant genetic effects overlapped; 24% of genes under local effects were also under distant effects, highlighting the polygenic nature of expression variation. We had previously used the haploid component of the same approach to identify segregating expression variation that was consistent with underlying single major effect loci in the same individual we studied here (Verta *et al.*, 2013). Most of the major genetic effects on expression variation was re-discovered in this study, along with many more genetic effects with typically smaller effects on expression variation (Supplementary text), demonstrating the high reproducibility of the approach for studying the genetic bases of expression variation.

### Effect of linkage versus allele specificity

We observed that local expression variation was associated with a complex mix of *cis*, *trans* and compensatory *cis*-*trans* effects. Most local effects acted in *cis* (53%), consistent with earlier studies (Ronald *et al.*, 2005, Pickrell *et al.*, 2010, Cheung *et al.*, 2010, Zhang *et al.*, 2011, Gruber *et al.*, 2012, Massouras *et al.*, 2012, Meiklejohn *et al.*, 2013, Lagarrigue *et al.*, 2013, Battle *et al.*, 2014). These observations are in line with the view that a significant part of local effects represent allelic sequence or chromatin variation that alter the rate of transcription or mRNA accumulation in general (Wittkopp and Kalay, 2012, Shlyueva *et al.*, 2014), but see (Gerstein *et al.*, 2012). However, our results indicate that a non-negligible portion (up to 12% of tested) of local variation acted in *trans*. We observed that the proportion of local effects acting in *trans* increased to a third when we investigated local effects that had effects across all diploid genotype combinations (because some local *cis* effects were not observed in homozygotes due to compensatory *trans* effects, Fig. 4E). Although rarely dissected in a systematic fashion, indications that *trans* acting effects might represent a significant portion of local variation have been reported (Ronald *et al.*, 2005, Cheung *et al.*, 2010, Zhang *et al.*, 2011, Lagarrigue *et al.*, 2013). Taking into account the resolution of our study (66 segregant progeny), the factors associated with local *trans* regulation could be independent of the focal gene but situated on the same chromosome. It is also possible that some local *trans* effects influence self-regulation of transcription, such as in the well-documented case of the yeast gene *AMN1,* where the gene product regulates the transcription of its locus (Ronald *et al.*, 2005). Further fine-mapping would help to determine whether the local *trans* effects identified here were caused by variation in the gene itself, or independent genes in linkage.

A complex architecture likely underlies most local effects. In a considerable portion (39% of tested) of the local *cis* effects, expression was allele-specific in heterozygotes, but no difference between homozygous genotypes was observed (Fig. 4E). This pattern is most consistent with compensation between *cis* and *trans* effects; focal effects are cancelled in homozygotes but not in heterozygotes where the compensatory factors cancel each other (Fig. 4F, Landry *et al.*, 2005). In addition, 165 (18% of tested) focal genes showed patterns consistent with the compensatory effect being stronger than the focal effect, leading to a change in signs between homo- and heterozygotes (Fig. 4F, Goncalves *et al.*, 2012). Our results show that the putative compensatory factors must be in very close linkage to the focal gene because their effects are observed in a segregating progeny.

Overall, compensation seems to be a common feature of transcriptional networks and suggests that expression variation is (or has been) under stabilizing selection (Wittkopp *et al.*, 2004, Landry *et al.*, 2005, Tirosh *et al.*, 2009, McManus *et al.*, 2010, Goncalves *et al.*, 2012). Stabilizing selection allows mutation of small effects that increase and decrease expression levels in opposing directions to accumulate in populations. Recombination between compensatory factors may give rise to combination of genotypes that show extreme phenotypes, which represent a form of genetic load (see: Charlesworth 2013). Although compensation seems to be widespread across organisms, it has remained unknown if the compensatory factors were dispersed over chromosomes or not because earlier studies have not measured linkage. Close linkage between the compensatory and focal effects such as seen here could be favored by natural selection as recombination would tend to break compensatory associations, thus unleashing negative expression variation or leading to other maladaptive expression traits. The means by which closely linked compensatory effects evolve remains to be determined, but could involve either preferential establishment of compensatory effects near the focal effect locus, or “migration” of effects to close linkage by means of genomic rearrangements (see also: Yeaman, 2013).

### Evolution of expression variation

Previous studies have established that expression divergence due to *cis* variation is more common between species, compared to the more widespread *trans* variation observed within species (Wittkopp *et al.*, 2008, Emerson *et al.*, 2010, Schaefke *et al.*, 2013). These patterns are likely influenced by dominance relationships between the underlying allelic variants; populations may preferentially accumulate recessive effects because they are less likely to reach fixation (Lemos *et al.*, 2008). Consistently, *trans* variation seems to be much more recessive than *cis* variation (this study, Lemos *et al.*, 2008, McManus *et al.*, 2010, Emerson *et al.*, 2010, Gruber *et al.*, 2012, Schaefke *et al.*, 2013). However, because previous studies have not distinguished between local/distant and *cis*/*trans* effects (but see Gruber *et al.*, 2012), it has remained unclear whether effect linkage or allele-specificity underlie these differences in inheritance.

Our results indicate that the level of dominance in expression phenotypes seem to be dependent on the allele specificity of genetic effects, more or less independently of their linkage to the focal gene. First, the majority of distant effects, all acting in *trans*, were dominant/recessive (Fig. 6B). Second, expression phenotypes under local *cis* effects were mostly additive (Fig. 6B). Third, a marked difference was observed in the inheritance of expression phenotypes under local *trans* effects versus those under *cis* effects. In a majority of local *trans* effects, heterozygote expression was not statistically different from one of the homozygotes, consistent with dominance by one of the alleles. The level of phenotypic masking, as measured by the D/A ratio, indicated that dominance was not complete (Fig. S8A). Previous studies indicate that such partial dominance is a common feature of expression variation (Gibson *et al.*, 2004, Lemos *et al.*, 2008). Some uncertainty remain in these results, as some statistically dominant/recessive effects had D/A ratios close to zero, indicating a conflict between the two measures for dominance (e.g. Fig. S8A). We favor interpreting the results based on statistical expression difference rather that the D/A ratio because the D/A ratio does not take into account variance within genotype classes and is thus more limited in terms of interpretation (Gibson *et al.*, 2004). Here also, further dissection of local expression variation with higher resolution would be informative.

Our results suggest that further refinement in the assumed evolutionary implications of local variation can be gained. Gruber *et al.* (Gruber *et al.*, 2012) proposed that due to their larger phenotypic effects and more additive nature in diploids, local (in majority *cis*) genetic variation are under more efficient natural selection and thus either reach fixation, or are eliminated quicker than distant (in majority *trans*) variation, leading to local effects being preferentially observed between species relative to within species (see also: Lemos *et al.*, 2008). Our results suggest that preferential fixation of effects, if dependent on the mode of inheritance, should not apply to all local variation but depend on their allele specificity.

### Nature of *cis* and *trans* effects

The patterns of inheritance reported in the present study are consistent with *trans*-acting variants influencing the focal gene via a diffusible molecule. In contrast to *cis* effects where each allele is presumably transcribed in an independent quantity, diploid focal genes under *trans* effects are exposed to both effect alleles as well as other factors influencing their transcription (Wittkopp *et al.*, 2004). In this context, we propose that the widespread dominance/recessivity of *trans* acting variation could emerge from the complex nature of gene transcription in a similar way as initially described by Sewall Wright in the form of the physiological theory of dominance (Wright, 1934), and later much elaborated by Kacser and Burns (1981) as the metabolic theory. According to this scenario, diffusible *trans* effects would influence expression through their complementary functions in association with the multitude of other proteins and RNAs involved in transcription. This might take place at chromosome locations containing regulatory sequences in the form of combinatory binding of proteins (Gerstein *et al.*, 2012), in the form of interactions within a transcription factor – co-factor complex (Slattery *et al.*, 2011), or interaction within a transcription factor protein complex, for example. Because most *trans* effects would represent non rate-limiting factors in gene transcription in this scenario, their variation would generally follow a relationship of diminishing returns where only the complete absence of a factor due to a homozygous genotype would lead to an observable expression difference. This haplosufficient characteristic was present in the majority *trans* effects, whether local or distant relative to the focal gene (Fig. 5).

Further, we show that distant *trans* effects fulfill a key prediction of the physiological and metabolic theories (Wright, 1934, Kacser and Burns, 1981, Phadnis and Fry, 2005), which is that dominance becomes stronger with larger effects (Fig. 6C). This characteristic follows from the nature of the reactions in question (in our case, gene transcription), which contain multiple interdependent steps (like the transcriptional protein complex and interacting enhancer or repressor proteins, for example). In the theory of Kacser and Burns (1981), these steps depend on one another because they share the same pool of substrates and products, which tends to buffer the system to variation in any one component. It is reasonable to assume that this could also apply to gene transcription not only in the form of substrates, but also as interaction states between different proteins. When the system contains many steps, the response between phenotype and genotype takes the form of a hyperbolic curve of diminishing returns (figure 4 in Kacser and Burns, 1981). The impact of mutation in any one of the constituents on the resulting phenotype will depend on their importance for the functioning of the whole system. Mutations in factors that have small effects on the total flux through a reaction when they are homozygous will be additive, while mutations in components that are essential for flux will be dominant/recessive (Fig. 6C, Kacser and Burns, 1981).

Overall, distant *trans* effects of small magnitude were more additive than those that explained a large proportion of expression variation, consistent with the prediction of the metabolic theory. Our results show that unlike *trans* effects, larger *cis* effects were more additive. They therefore seem to represent rate-limiting factors in gene transcription that could correspond to their structural instead of diffusible nature. By representing differences in the DNA sequence, *cis* effects might alter the overall affinity of the whole panoply of transcriptional regulators to the DNA strand and hence yield a largely additive effect in heterozygotes irrespective of their effect sizes.

In summary, our systematic dissection of the relationships between linkage, allele-specificity and dominance in expression variation represents a significant step towards a better understanding of the role of expression variation in past, present and future evolution. The approach developed in this report can be used to investigate expression divergence between and within a wide range of wild plant species. Haploid/diploid analyses akin to the one reported here are becoming increasingly feasible through the use of doubled haploid lines of angiosperms for example (Seymour *et al.*, 2012), allowing future investigation of the evolutionary modes discovered here in other phyla.

## Materials and Methods

A white spruce individual (known as 77111) growing at Cap Tourmente, Québec, Canada (+47° 4’ 1.28”, −70° 49’ 4.03”) was self-fertilized in May 2010 by Jean-Marc Montminy of Canadian Ministry of Natural Resources. Pollen of 77111 was obtained from Dr. Jean Beaulieu of Canadian Forest Service. Cones were collected on August 18^th^ 2010, and stored in paper bags in room temperature and protected from light until they were used for the study. The seeds were stratified for 4 wk and germination process was initialized as described in Verta *et al.* 2013 (empty seeds and seeds where embryos had obvious morphological anomalies were discarded, see Supplementary text). Seeds were dissected under a microscope. Megagametophyte and embryo tissues were flash-frozen on liquid nitrogen and stored in −80 degrees C until they were used for RNA extraction.

RNA extraction and quality control was carried out as described in Verta *et al.* 2013. Individual RNA-seq libraries were constructed from each mRNA sample with the TruSeq RNA sample preparation kit (Invitrogen), with minor modifications to the manufacturer’s protocol. The indexed sample libraries were combined to single lanes in equal amounts in batches of 24 samples for megagametophytes and 23 samples for embryos. A total of seven lanes of 100 bp Illumina HiSeq-2000 paired-end sequencing of the pooled libraries was carried out at the Genome Québec Innovation Centre in Montreal, Canada. One pooled sample was run per lane. One particular pool was sequenced on two lanes and the reads were combined. The number of reads per lane is given in table S1. Read quality was verified with the FastQC software (www.bioinformatics.babraham.ac.uk/projects/fastqc/) and Illumina sequencing adapters were trimmed from the reads using TrimGalore (www.bioinformatics.babraham.ac.uk/projects/trim_galore/).

### Alignment of RNA-seq reads

Reads from each individual sample were aligned to white spruce gene models representing single genes (Rigault *et al.*, 2011) using Bowtie 2 (Langmead and Salzberg 2012) in the local alignment mode with default parameters. The resulting .sam files were transformed into .bam format, sorted and likely PCR duplicates were removed with the SAMtools (Li *et al.*, 2009) functions *view, sort* and *duprem*, after which the files were indexed with the function *index*.

### Expression quantification per transcript

Per transcript read counts were generated using the HTSeq software (Anders *et al.*, 2014) with the *interception-strict* read counting mode. Reads having a mapping quality less than 30 were not considered. Tissue-preferential expression was analyzed in R (version 2.15, R Core Team, 2013) with the package ‘edgeR’ (Robinson *et al.*, 2010). Samples within in each tissue type were considered as biological replicates. Normalization factors, and common and tag-wise dispersion estimates were calculated with the edgeR functions *calcNormFactors(…method=*‘*TMM*’), *estimateGLMcommonDispersion* and *estimateGLMtagwiseDispersion*. Total library size for each sample was used in the calculation of the offset parameter, as is default in edgeR. Consequently, gene-wise expression differences between the haploid and diploid samples were summarized with the *glmLRT*.

### SNP discovery in megagametophytes

Transcribed SNPs were identified individually for each megagametophyte sample with the FreeBayes software (version 0.9.8, Marth, 2012), specifying ploidy as 1 and enabling the standard filters. The mapping quality, read coverage and segregation of each putative variant was then analyzed with custom Python (version 2.6) and Biopython (Cock *et al.*, 2009) scripts. A final set of SNPs were selected based on the following requirements: i) mean mapping score of reads overlapping the SNP position was higher than 30 in at least 90% of the samples where the variant was observed, ii) the nucleotide position of the variant was covered by at least one read in 90% of all the samples, iii) the frequency of the two alternative nucleotides were within the 95% IC of Mendelian segregation, i.e. between 25 and 41 observations when N=66, iv) at least 90% of the SNPs fulfilling the criteria (i-iii) for a given transcript were linked in 90% of the samples, i.e. if the transcript had SNPs that were not linked in 90% of the samples (which could be the case if reads from two similar transcripts map to the same reference sequence) none of the SNPs were considered for that gene. Reproducibility of base calls in the variant positions were investigated by comparing all aligned nucleotides overlapping a variant position within each sample.

### Embryo genotypes

Variants for embryo samples were called with FreeBayes, specifying ploidy as 2 and enabling the standard filters. The algorithm was given the genotype (specified as the .vcf file) of the associated megagametophyte sample as *a priori* information (option *-variant-input*). Custom Python scripts were then used to extract the genotype and the read coverage of each segregating SNP, as identified in the megagametophytes. Self-fertilization was validated by comparing embryo genotypes over the complete set of segregating SNPs against megagametophytes from the same seed.

### Association testing in megagametophytes

**Local effects** - Each gene was identified as one of the two maternal genotypes based on transcribed SNPs. Differential expression was tested considering samples as biological replicates of one of the two alleles and testing expression difference between the two groups. Two complementary approaches were used. First, a single SNP was used to assign samples to alleles. Then, gene-wise counts generated with HTSeq were used in the testing of expression differences between alleles with the R package edgeR as described above concerning tissue-preferential expression. Normalization factors, dispersions and offsets were calculated based on all reads in each sample. *Q-*values were calculated from *P*-values using the R package ‘qvalue’ (Storey, 2002).

In the second approach, gene expression level for each gene/sample was represented as the sum of read coverages over all heterozygous nucleotide positions (one if the gene had only one segregating SNP, or many if the gene contained a set of linked heterozygous SNPs). A minimum distance of 100 bp between SNPs within a given gene was required, corresponding to typical read length. Expression differences between allele classes were tested as above using edgeR. Offset parameter for the edgeR model was calculated sample-wise as the logarithm of the sums of alternative and reference allele read counts observed for each sample.

A bias towards higher expression of the allele represented by the reference has been reported in numerous studies where reads from a heterozygous individual are mapped to a reference (Stevenson *et al.*, 2013). Our approach takes this possible source of bias into account in two ways. First, read counts are summarized over linked SNPs within a gene, and because the two alleles (which are unrelated to the genotypes used in construction of the white spruce transcriptome reference) are anticipated to contain alternative and reference nucleotides in an equally frequency, no bias is expected. Second, we took into account the total number of reads overlapping the alternative and reference nucleotides, and used this as a baseline (the offset parameter) with which the abundances of the expression level of each allele were corrected. In the cases where only one allele was different for only a single SNP with respect of the reference, higher expression level was observed equally in the alleles represented by the alternative and reference nucleotides (121 and 108 cases, respectively).

**Distant effects** - Expression level of the focal gene was represented by the sum of reads overlapping the gene model, as the focal gene genotype was considered to have no impact in the case of distant effects. First, low-resolution linkage blocks were defined with the R package ‘qtl’ (Broman *et al.*, 2003), using the function *formLinkageGroups* with the options *max.rf=0.35* and *min.lod=6*. Genotypes of all other genes were initially tested against focal gene expression in the cases where both the focal gene, and the variant against which it was tested were covered by at least one read in more than 90% of the samples. With these results in hand, we determined the genetic distance over which 95% of the association signals from local effects were detectable on other loci (30 cM, *q*-threshold>0.01, Fig. S9). Accordingly, the final set of tested distant loci was filtered to include only loci whose genetic distance to the focal locus was at least 30 cM.

### Association testing in embryos

***Cis* effects** - Testing for *cis* effects was performed for genes in which samples heterozygous for the two focal alleles were observed in a Mendelian frequency, i.e. between 25 and 41 samples when N=66. Expression level for each allele was represented as sums of read counts over all heterozygous, linked variants as in the megagametophytes. Each allele in each sample were considered as replicate measures, and the difference between allele classes was tested with edgeR as described for the megagametophytes. This procedure takes into account sample-wise variation in total library size and potential reference mapping bias as described for the megagametophytes.

**Effects in homozygous diploids** - Embryo genotypes were called from transcribed SNPs with custom Python scripts, taking into account the genotype of the associated megagametophyte. Testing for differential expression between homozygous genotypes was performed for genes in which the observed genotypes segregated in a Mendelian 1:2:1 ratio, that is, heterozygotes were observed in a frequency from 25 to 41 and homozygotes in a frequency from 10 to 24. Testing procedure was identical to that described above concerning the megagametophytes, i.e. the edgeR package was used to test gene-wise counts and normalization factors, dispersions and offsets were calculated based on all aligned reads.

**Inheritance –** An edgeR model for gene-wise expression levels in focal gene according to local or distant genotypes was constructed as described above in the case of homozygote difference. Contrasts between heterozygous genotype versus the two homozygous genotypes were tested with the *makeContrasts* function of edgeR. Significant contrast between heterozygotes and both homozygous classes was designated as evidence for additivity. A single significant contrast was designated as evidence for dominance of the non-significant genotype. If neither contrast were significant the inheritance of the distant effect was designated as “unknown”. For distant effects, we identified a non-redundant set of association pairs by selecting one additive or dominant/recessive association per linkage block per focal gene (lowest *P-*value selected).

**Effect size** - Strength of association in likelihood-based models can be measured by a coefficient of correlation based on the likelihood ratio, the R*_LR_*^2^ (Supplemental text, Magee, 1990, Nagelkerke, 1991). Similar to the standard coefficient of determination in association studies (e.g. Laurie *et al.*, 2004), this parameter can be broadly interpreted as the proportion of explained variation in the phenotype accounted by a genetic effect (Sun *et al.*, 2010). Use of R*_LR_*^2^ as a measure of effect size follows from the fact that allelic variation in a genetic variant that is strongly associated with expression variation changes strongly the expression level (Rockman and Kruglyak, 2006). Consistent with this, high values of R*_LR_*^2^ corresponded to genetic effects that were associated with single major effect loci and median R*_LR_*^2^ closely corresponded to the median effect size previously estimated for eQTL in a yeast cross (Supplemental text, Brem and Kruglyak, 2005).

R*_LR_*^2^ was calculated from the likelihood ratio following (Magee, 1990);

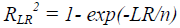

where *LR* is the likelihood ratio of the model (provided by the *glmLRT* function of edgeR) and *n* is the sample size. In haploids, this model included all samples assigned to one or the other allele. In diploids, the model included only homozygous genotypes.

**D/A ratio** - D/A ratio was calculated as defined in Gibson *et al.*, (2004). Relationship between R*_LR_*^2^ and the D/A ratio was tested with the R base function *lm*, considering D/A ratio as the response variable and R*_LR_*^2^ as the linear predictor.

## Data access

Scripts used for the analysis, variant calls (.vcf files) and gene expression tables are available from the Dryad database (Dryad allows data submission only following manuscript acceptance). Raw sequence data is available through the NCBI Sequence Read Archive (submission ongoing).

## Acknowledgements

We would like to thank Dr. Jean Beaulieu, Marie Deslauriers of Canadian Forest Service as well as Anne Savary and Jean-Marc Montminy of Quebec Ministry of Natural Resources for providing self-fertilized seed. We would also like to thank Dr. Brian Boyle for guidance in RNA-seq library preparation as well as Guillaume Diss and Jean-Baptise Leducq for helpful comments on the manuscript and the MacKay and Landry labs for support and insightful conversations. Funding to JM was from Genome Canada and Genome Québec as a part of the SMarTForests project and from an NSERC Discovery grant. Funding to CRL in ecological genomics comes from a NSERC Discovery grant and a salary fellowship from CIHR.

