## Supplementary Materials for "The Genetic Landscape of Transcriptional Networks in a Combined Haploid/Diploid Plant System"

### Supplemental Text

#### Distant effect hotspots

We compared the positions of distant effects to investigate whether the underlying loci were distributed randomly among the linkage blocks. Because most distant effects that influenced the same focal gene and mapped to the same linkage block were likely diagnostic of the same association, we reduced the set of distant effect loci to one per focal gene per linkage block, keeping the association with the lowest *P*-value. We then compared the number of non-redundant distant effects per linkage block to the number of effects that was expected if the effects were distributed randomly across all blocks, taking into account the number of tested SNPs in each block. Blocks 9-12 showed the highest difference between observed and expected numbers of effects, ranging from 56% to 77% above the expected, although the differences were not statistically significant ($P<0.24$).

We also analyzed the number of focal genes associated per individual distant effect locus. The single distant locus that had the most widespread effect was associated with the expression level of 543 focal genes. These included 250 mapped focal genes and 293 non-mapped focal genes. The distant effect locus (GCAT: GQ03105_C20.1, GenBank: BT107266) corresponded to a gene that encodes for a putative pseudouridine synthetase (Pfam: PF01416). This gene could not be assigned to any of the 12 larger linkage blocks, but appeared unlinked from all other analyzed genes (likely a consequence of segregation distortion Verta J.-P., Landry C.R. & MacKay J., unpublished data). Therefore, its linkage to other distant effect loci could not explain the number of focal genes associated with genetic variation in this locus.

#### Magnitudes of local and distant effects

Expression variation, as any other complex phenotype, is influenced by multiple loci, most of which have small effects (Brem *et al*., 2002, Brem and Kruglyak, 2005). We hypothesized that this was also the case for local and distant effects identified in the haploids. The approach outlined in the main text models genetic effects on gene expression as likelihood functions between focal gene expression levels (read counts) and genotypes. Therefore, the proportion of expression variation that could be accounted for by the associated genetic effects could not be summarized with an $R^2$ parameter as is the usual case in least squares regression-based analyses. Instead, we calculated the likelihood ratio based $R_{LR}^2$ in order to quantify the improvement in the likelihood model in explaining expression variation, versus a null model with only the intercept parameter (Magee, 1990, Nagelkerke, 1991, Sun *et al.*, 2010). The median of $R_{LR}^2$, which can be broadly interpreted as the proportion of explained expression variation (Sun *et al*., 2010), was very close to the proportion of explained variation typically expected for eQTL (Fig. S4, Brem and

Kruglyak, 2005). We then compared the $R_{LR}^2$ of local and distant effects to the cases where expression variation between the alleles was described in previous work involving the same individual tree (Verta *et al.*, 2013). Our previous analyses indicated that these expression differences had relatively simple genetic basis with a single major causal locus that explained a large portion of expression variation in each case. Consistent with this, $R_{LR}^2$ of the majority of local and distant effects identified in this report were smaller than the cases where a single major variant was previously observed to underlie the segregating expression difference (Fig. S4, $P<2\cdot10^{-16}$). This result suggested that the likelihood models captured associations where expression variation was caused not only by a single major genetic effect, but also cases where the associated variant had a small effect on expression variation and likely represented part of a polygenic variation.

Overall, the distribution of $R_{LR}^2$ values of all distant effects was shifted toward higher values than those of local effects. In fact, no $R_{LR}^2$ value inferior to 0.2 was observed in the case of distant effects. We hypothesize that this may have been a consequence of a higher multiple testing constraint and the fact that $R_{LR}^2$ values were calculated based on likelihood ratios of the identified associations. The associations that yielded only small increased likelihood in comparison to the null model may have not reached our *q*-value threshold.

#### Gene-wise versus SNP-wise approaches

In order to facilitate comparison of allele-specific expression levels across haploids and diploids, we defined a set of local effects that could be identified based on reads overlapping transcribed SNPs. Testing of *cis* effects in diploids leverages on counting the number of RNA-seq reads overlapping heterozygous SNPs that can be used to distinguish between the two alleles of a heterozygous genotype (referred to as SNP-wise approach, Fontanillas *et al.*, 2010). Reads not containing SNPs are non-informative towards *cis* effects in diploids, and the method is therefore inherently less powerful to find expression differences than a haploid approach where all reads overlapping an allele can be used to estimate expression levels (referred to as gene-wise approach). Our system greatly facilitates the study of *cis* effects based on SNP-wise counts because identification of multiple phased SNPs in haploid tissue of heterozygous individual is relatively straight forward, and allows counts over multiple variant positions to be summarized within single genes (Methods).

We compared local effects identified in the haploids with the two complementary approaches, i.e. based on gene-wise and SNP-wise read counts. Nearly all of the local effects (84%, $q<0.01$) identified with gene-wise approach were observed with the SNP-wise approach also. The magnitudes of the expression differences between alleles were highly correlated in the two cases (Spearman $r=0.96$). Overall, a smaller frequency of local associations (36%) was observed compared to the case where all reads for a given allele were included in the expression estimate (48%), as was anticipated. The power and reproducibility of the SNP-wise approach relative to gene-wise approach was dependent on the number of heterozygous positions available for generating the expression estimate. With the $q$-value threshold of 0.01, estimates based on single SNPs gave a lower frequency of statistically significant effects (24%) but a similar frequency of concordant effects (83%) than estimates based on more than one SNP (44% and 84%, respectively).

#### Test of seed viability

White spruce is normally outcrossing, and self-fertilization is expected to lead to a lower than normal level of seed viability. We tested seed viability by sowing 1755 stratified seeds in October 2010 (germinated in growth chamber). After one month, 210 seeds had germinated (12%) and after three years 166 (9.5%) had survived. We sowed 225 germinated seeds again in July 2012 (germinated in greenhouse), which had similar germination percentage (25 plants).

#### Supplemental Figures and Tables

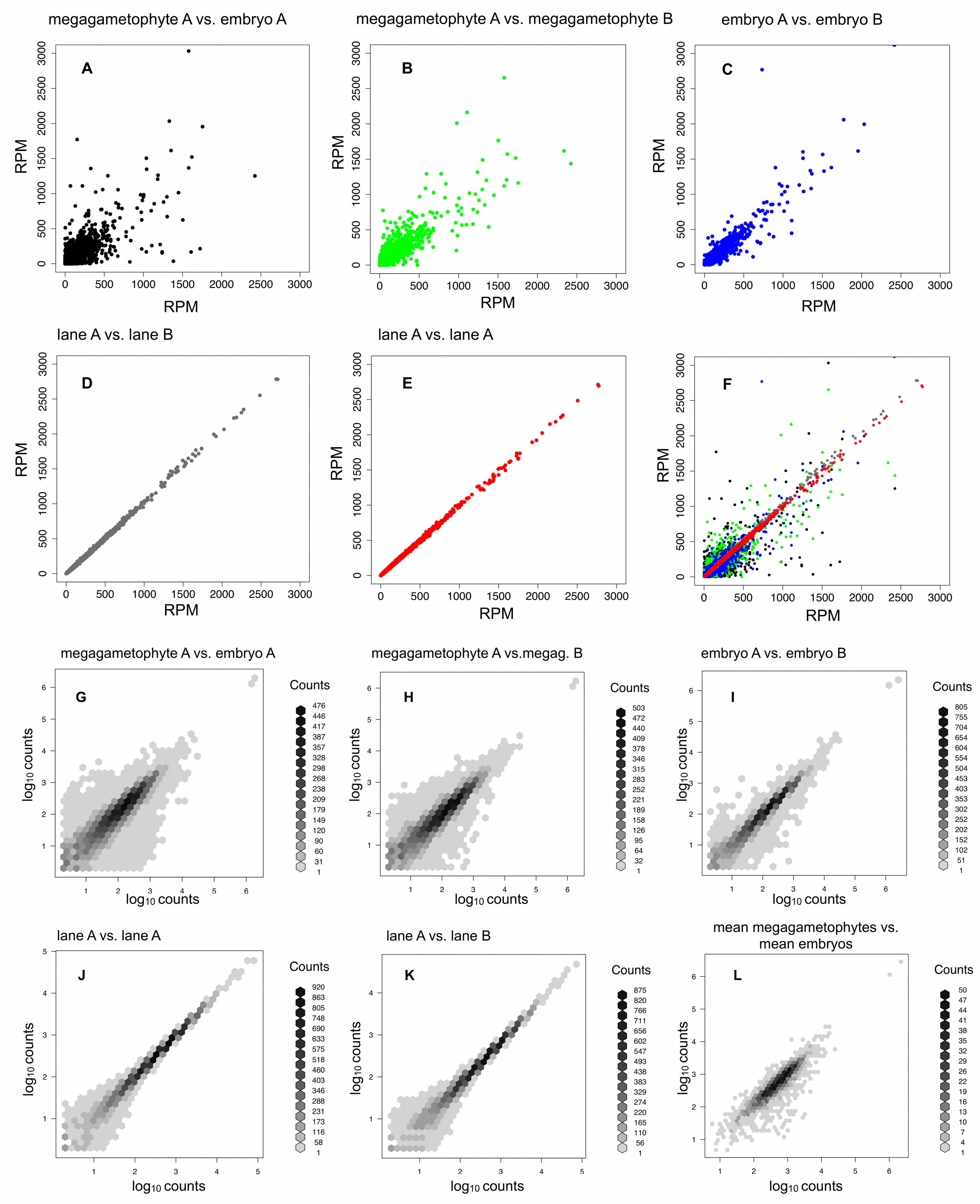


**Figure S1. Comparison of RPM- and $log_{10}$-normalized counts across sample pairs.** A-F: RPM (Reads Per Million) normalization takes into account the differences in library size between samples and is used here only for illustration purposes. Pairs represent individual samples. **A)** black points; randomly selected megagametophyte and embryo of the same seed, **B)** green points; two randomly selected megagametophytes, **C)** blue points; two randomly selected embryos, **D)** grey points; sample run twice on the same lane with different index, **E**) red points; sample run on two lanes, **F)** all previous comparisons plotted at once. G-L: $Log_{10}$ –normalized counts (genes containing zero counts are omitted). Normalization does not take into account differences in library size and is used here only for illustration purposes. **G-K)** comparisons as above for RPM counts, **L)** mean counts of all megagametophyte and embryo samples.

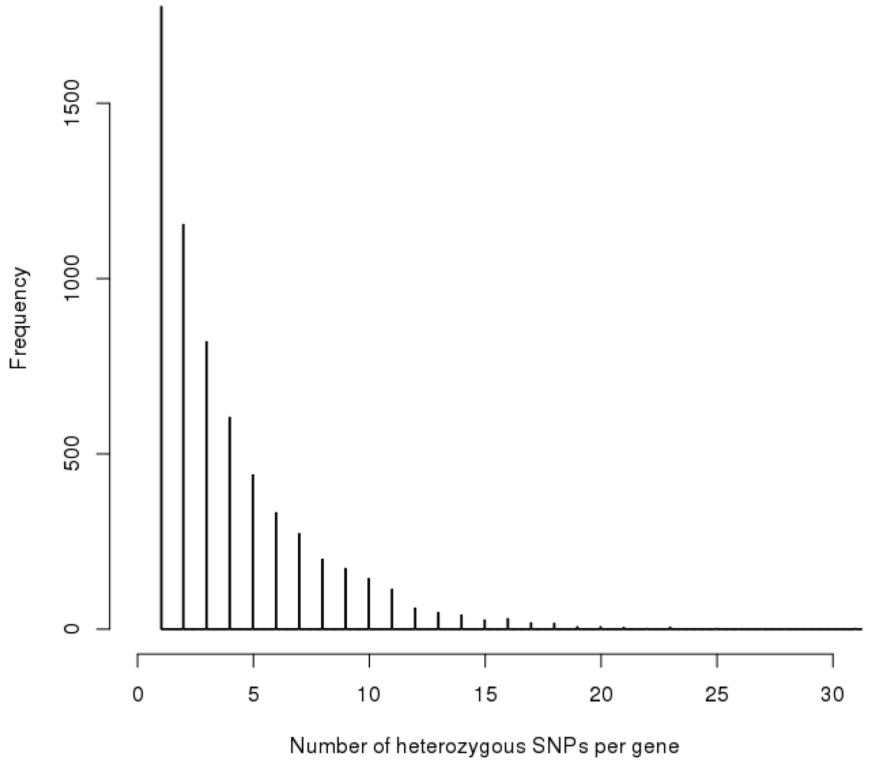


**Figure S2. The number of heterozygous SNPs per gene.**

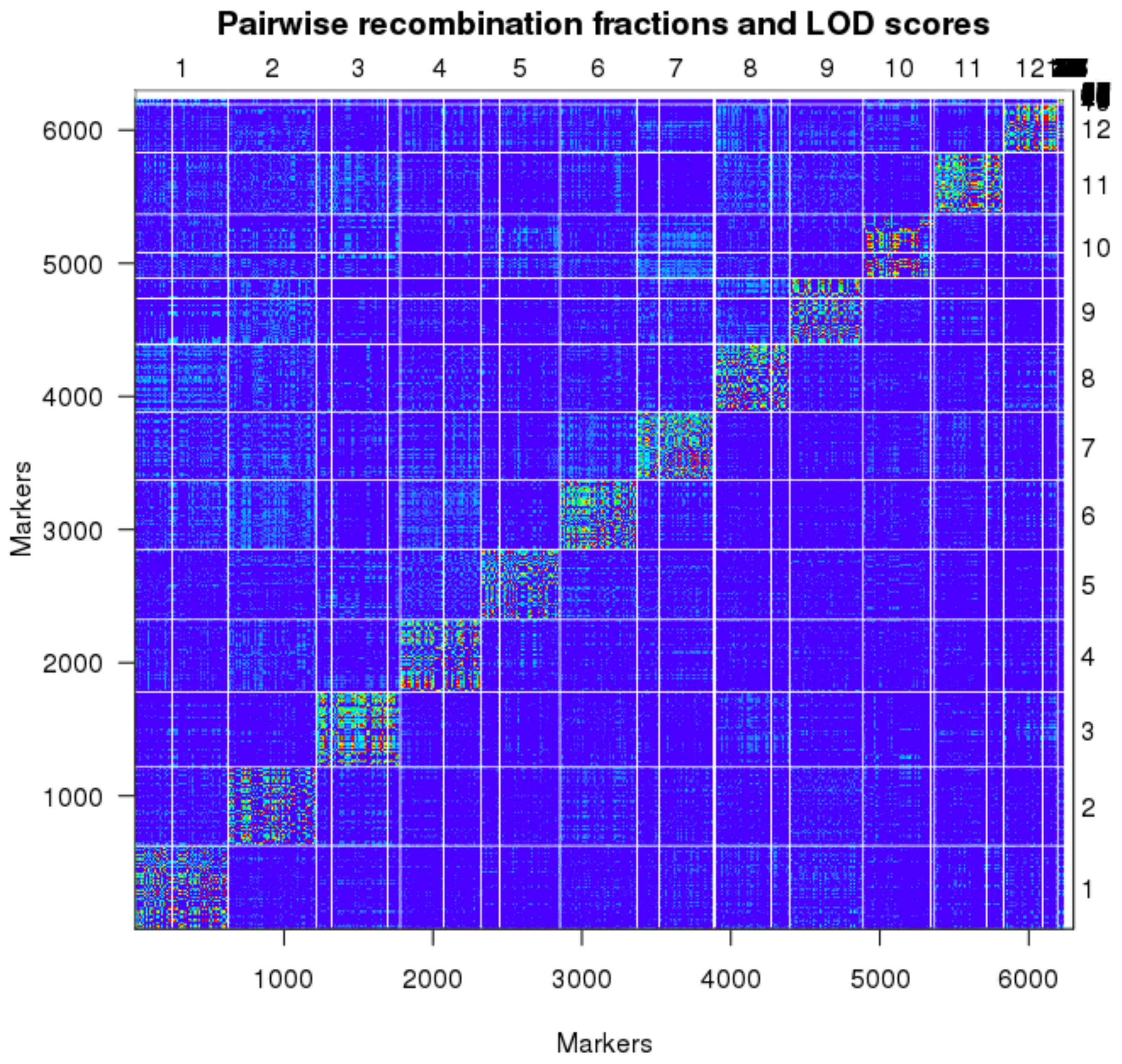


**Figure S3. Linkage groups.** Upper and lower triangles show pairwise recombination fractions and LOD-scores, respectively. Values are depicted as colors from blue (inferior values) to red (superior values). Right y-axis and upper x-axis show linkage groups.

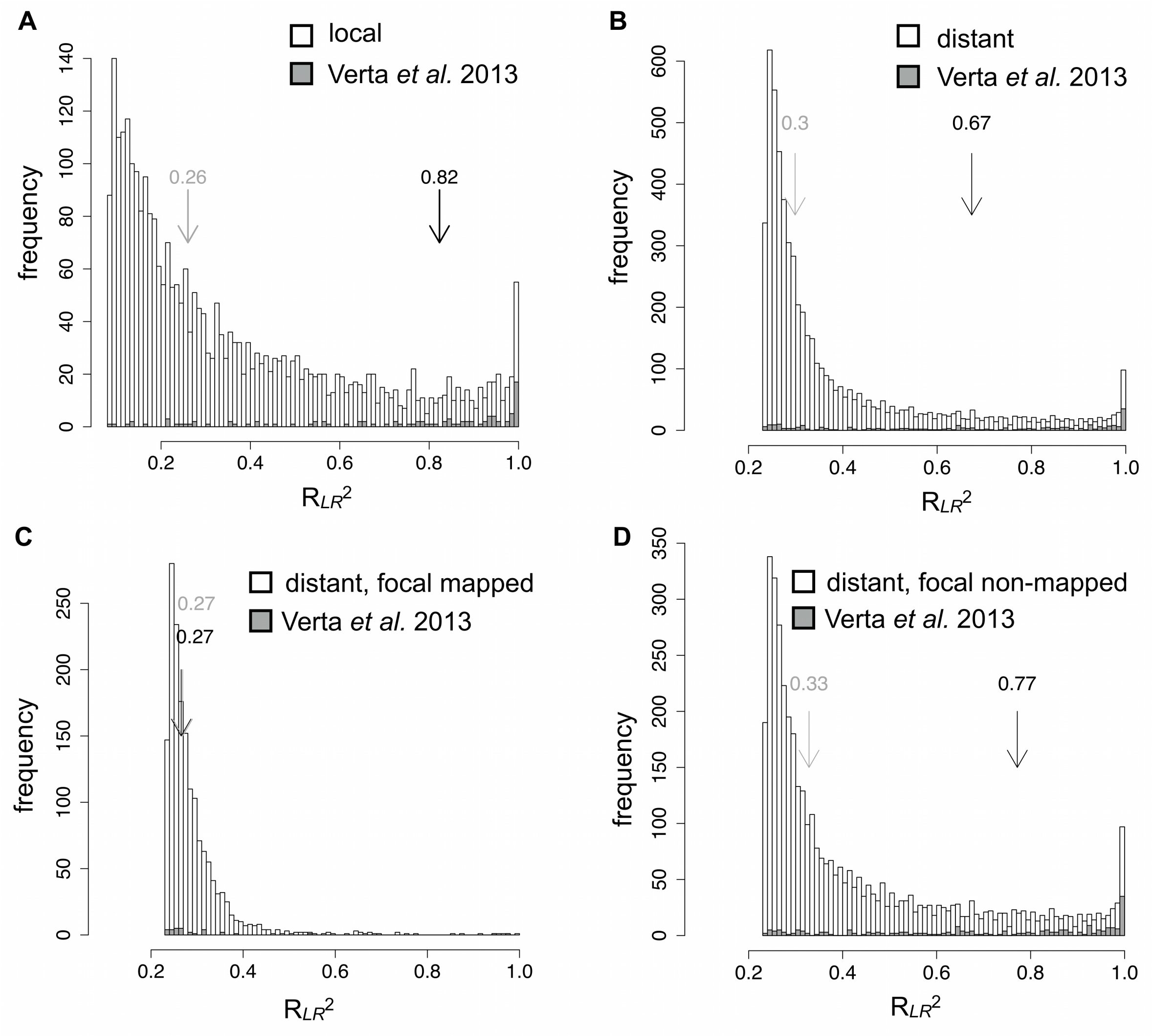


**Figure S4. Magnitudes of genetic effects on gene expression in haploids.** Most discovered associations between expression variation and local/distant variants were described by likelihood models that had lower goodness-of-fit versus the genes where expression variation was associated with single major effect loci discovered in a previous study (Verta *et al*., 2013), suggesting that most associated variants represented a part of polygenic variation. Relative gain in explaining power of likelihood models of local and distant variants versus a null model is described by $R_{LR}^2$ (Nagelkerke, 1991). Arrows depict medians of all effects (gray) and the subset of effects identified as due to single major effect loci in previous study (Verta *et al*., 2013, black). **A)** $R_{LR}^2$ of local variants. **B)** $R_{LR}^2$ of all distant variants. The $R_{LR}^2$ of the associated distant locus with the lowest P-value is reported in cases where the focal gene was associated with multiple distant loci on the same linkage block. **C)** $R_{LR}^2$ of distant effects influencing mapped focal genes. **D)** $R_{LR}^2$ of distant effects influencing non-mapped focal genes.

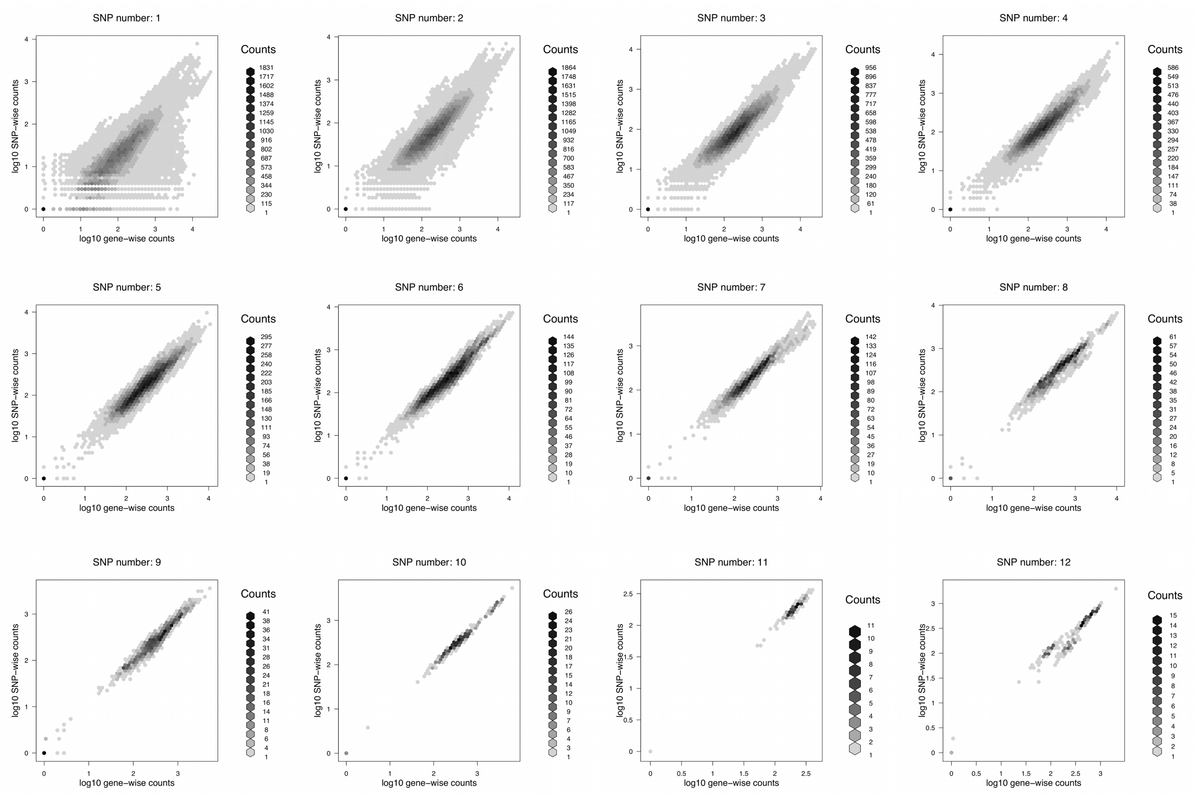


**Figure S5. Gene-wise versus SNP-wise counts per gene in the haploids.** Each plot represent $\log_{10}$ -normalized gene-wise counts (x-axis) plotted against SNP-wise counts (y-axis) of the same genes across all haploid samples. Number of SNPs in each case is given above each plot. Cases where 1 to 12 SNPs were used to estimate allele expression levels are given (the observed maximum number of SNPs per gene was 43).

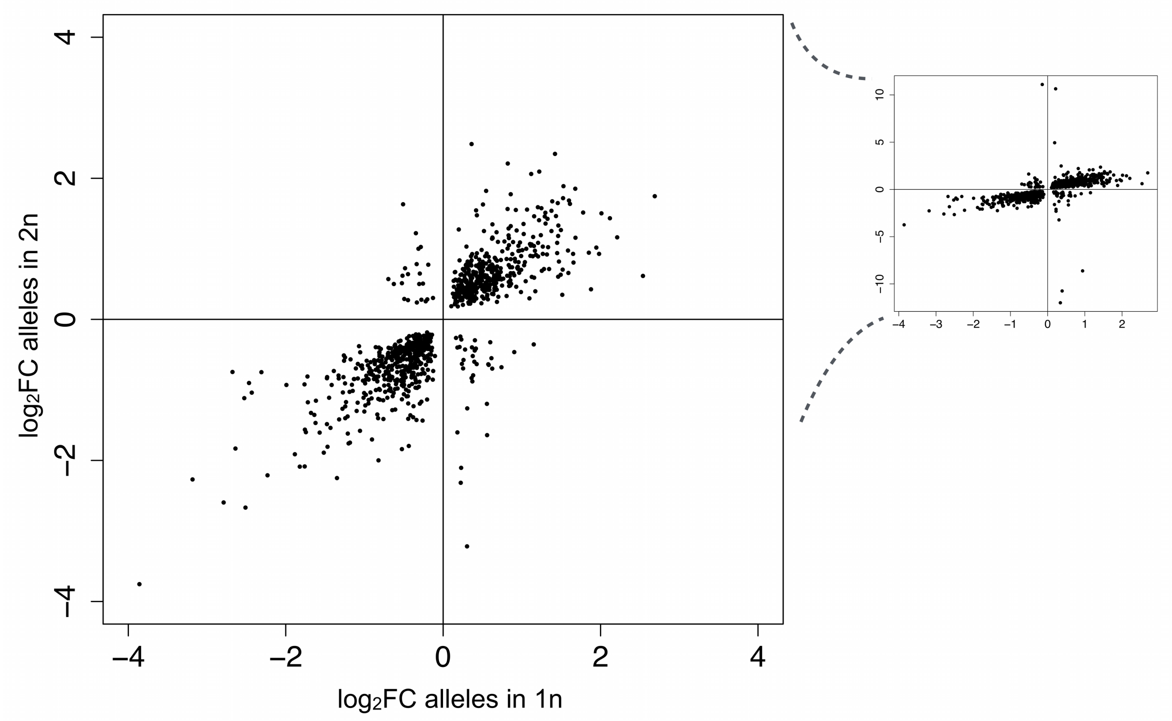


**Figure S6. Allele expression differences in genes under local *cis* effects.** Expression difference in haploids (x-axis) is plotted against expression difference between the alleles in heterozygous diploids.

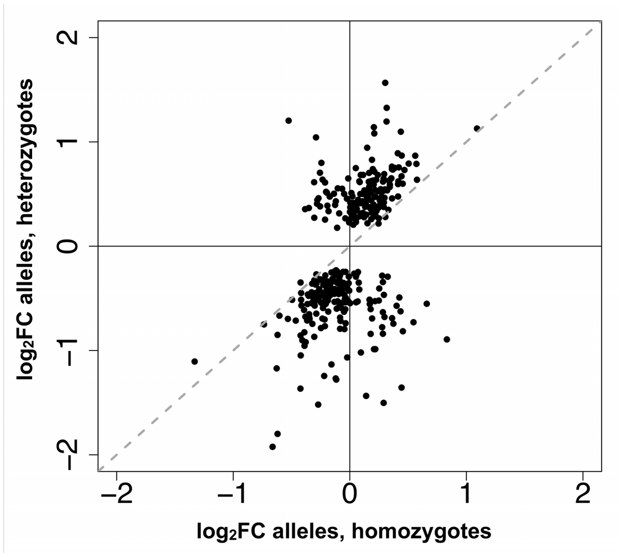


**Figure S7. Comparison of allele expression difference in homo- and heterozygous diploids in the cases where *cis* effects were observed in the absence of homozygote differences.** Diagonal dashed line marks equal difference between allele expression levels in the two cases.

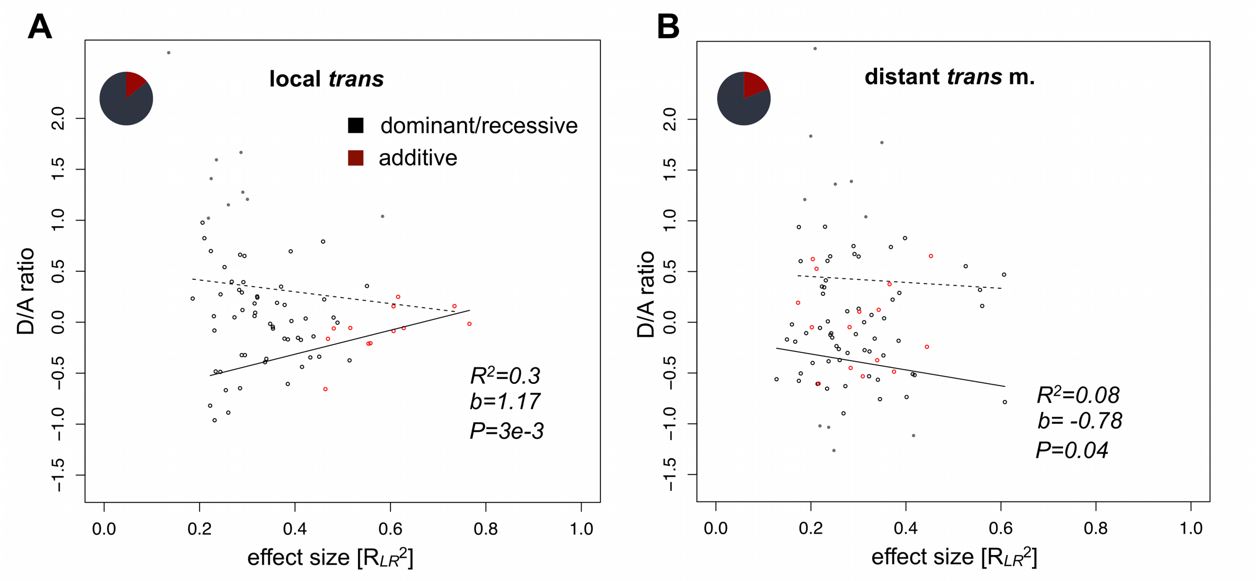


**Figure S8. Phenotypic masking by dominance versus effect size. A)** Local *trans* effects. **B)** Distant effect influencing mapped focal genes. Grey points depict under- or over dominant effects (not included in regression analysis). Lines represent linear model fit between D/A ratios (independently for negative and positive dominance) and strength of association are given for each category (non-significant regression represented by dashed line). Pie-charts show the proportions of additive (red) and dominant/recessive (black) effects in each category.

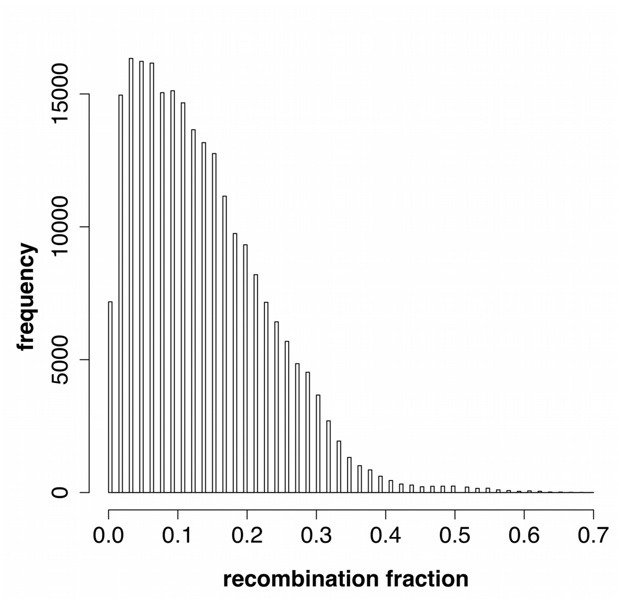


**Figure S9. Association signal from local effect on adjacent loci.** The number of adjacent loci that recapitulated a local association in the focal gene is showed in function of recombination fraction between the locus and the focal gene.

**Table S1. RNA-seq read yield per sequencing lane.**

| Pool # | # Samples | # Reads | # Bases |
|---|---|---|---|
| Embryo 1 | 23 | 214,803,221 | 42,960,644,200 |
| Embryo 2 | 23 | 81,036,043 | 16,207,208,600 |
| Embryo 2 | 23 | 135,396,216 | 27,079,243,200 |
| Embryo 3 | 23 | 214,732,094 | 42,946,418,800 |
| Mgg 1 | 24 | 207,479,401 | 41,495,880,200 |
| Mgg 2 | 24 | 212,830,751 | 42,566,150,200 |
| Mgg 3 | 24 | 246,596,517 | 49,319,303,400 |

**Table S2. Number of focal genes under local and distant effects at *q*-value < 0.01.**

| | local | distant | distant focal mapped | distant focal non-mapped |
|---|---|---|---|---|
| Number of focal genes | 3004 | 5770 | 1640 | 4130 |
| Tested genes | 6281 | 15051 | 6281 | 8751 |
| Percentage | 48% | 38% | 26% | 47% |

**Table S3. Local effects according to their allele-specificity and inheritance.** All reported effects influence non-preferentially expressed genes. Reported number represents focal genes where signs of expression difference between alleles in different genotypes did not change. Number of tested genes can change if they do not fulfill selection criteria for segregation frequencies in each case, are preferentially expressed in one tissue, or some genotypes are not expressed, or are expressed in less than 90% of the considered samples. The first row indicated which genotypes were tested and in which these criteria therefore needed to be fulfilled.

| Comparison | *a/A (1n) vs. aA (2n)* | | | *aa vs. aA vs. AA* | | | | | |
|---|---|---|---|---|---|---|---|---|---|
| Effect | *cis* | *cis* local | local *cis* | local *trans* | local homoz. difference | local *cis* additive | local *cis* d.r. | local *trans* additive | local *trans* d.r. |
| Number of focal genes | 1379 | 897 | 897 | 134 | 518 | 294 | 61 | 13 | 79 |
| Tested genes | 4429 | 1193 | 1692 | 1158 | 1200 | 356 | 356 | 133 | 133 |
| Percentage | 31% | 75% | 53% | 12% | 43% | 83% | 17% | 10% | 59% |

**Table S4. Distant effects according to their allele-specificity and inheritance.** Table represents non-redundant association pairs (total number of paired tests is given in parentheses) involving non-preferentially expressed genes that segregate in Mendelian frequencies.

| Comparison | *bb vs. BB* | | *bb vs. bB vs. BB* | | | |
|---|---|---|---|---|---|---|
| Effect | distant homoz. difference | | distant *trans* additive | | distant *trans* d.r. | |
| Focal gene | mapped | non-mapped | mapped | non-mapped | mapped | non-mapped |
| Number of focal genes | 109 (1171) | 1107 (34079) | 16 (542) | 94 (17393) | 70 (413) | 800 (7632) |
| Tested genes | 697 (5980) | 2460 (89800) | 86 (955) | 918 (25026) | 86 (955) | 918 (25026) |
| Percentage | 17% | 45% | 19% | 10% | 81% | 87% |
